# Post-vaccine epidemiology of serotype 3 pneumococci identifies transformation inhibition through prophage-driven alteration of a non-coding RNA

**DOI:** 10.1101/2022.09.21.508813

**Authors:** Min Jung Kwun, Alexandru V. Ion, Hsueh-Chien Cheng, Joshua C. D’Aeth, Sam Dougan, Marco R. Oggioni, David A. Goulding, Stephen D. Bentley, Nicholas J. Croucher

**Author notes:** These authors contributed equally (Min Jung Kwun, Alexandru V. Ion, Hsueh-Chien Cheng).

## Abstract

The respiratory pathogen *Streptococcus pneumoniae* (the pneumococcus) is a genetically diverse bacterium associated with over 100 immunologically-distinct polysaccharide capsules (serotypes). Polysaccharide conjugate vaccines (PCVs) have successfully eliminated multiple targeted serotypes, yet the mucoid serotype 3 has persisted despite its inclusion in PCV13. This capsule type is predominantly associated with a single globally-disseminated strain, GPSC12 (CC180), which was split into clades by a genomic analysis. Clade I, the most common, rarely underwent transformation, but was typically infected with the prophage ϕOXC141. Prior to the introduction of PCV13, this clade’s composition shifted towards a ϕOXC141-negative subpopulation in a systematically-sampled UK collection. In the post-PCV era, more rapidly-recombining non-Clade I isolates, also ϕOXC141-negative, have risen in prevalence. The low *in vitro* transformation efficiency of a Clade I isolate could not be fully explained by the ∼100-fold reduction attributable to the serotype 3 capsule. Accordingly, prophage ϕOXC141 was found to modify csRNA3, a non-coding RNA that inhibits the induction of transformation. This alteration was identified in ∼30% of all pneumococci, and was particularly common in the unusually-clonal serotype 1 GPSC2 strain. RNA-seq and quantitative reverse transcriptase PCR data demonstrated the altered csRNA3 was more effective at inhibiting production of the competence stimulating peptide pheromone. This interference with the quorum sensing needed to induce competence lowered the rate of spontaneous transformation, reducing the risk of the prophage being deleted by homologous recombination. Hence the selfish prophage-driven alteration of a regulatory RNA limits cell-cell communication and horizontal gene transfer, complicating the interpretation of post-vaccine population dynamics.

---

*Streptococcus pneumoniae* (the pneumococcus) is a globally-endemic gram-positive nasopharyngeal commensal bacterium and respiratory pathogen that causes both common (e.g. conjunctivitis, otitis media) and invasive (e.g. sepsis and meningitis) infections [1]. The species is genetically diverse, and subdivided into hundreds of strains, termed Global Pneumococcal Sequencing Clusters (GPSCs) by the Global Pneumococcal Sequencing (GPS) project [2, 3]. These strains encode particular combinations of accessory loci from the pneumococcus’ extensive pangenome [4], which can be exchanged between isolates through two classes of mechanism. The first is the movement of mobile genetic elements (MGEs), commonly represented by prophage, phage-related chromosomal islands (PRCIs) and integrative and conjugative elements (ICEs) in pneumococci [4]. The second is cell-driven transformation, which integrates exogenous DNA into the chromosome through homologous recombination [5]. This favours deletion of accessory loci, including MGEs, over insertion [6]. Therefore the two types of mechanism can be viewed as making conflicting contributions to bacterial evolution [7].

One of the most variable loci in the pneumococcal genome is the capsule polysaccharide synthesis (*cps*) locus, which encodes the machinery for generating the polysaccharide capsule, and therefore determines an isolate’s serotype [8]. Over 100 immunologically-distinct serotypes are expressed by pneumococci [9], which are associated with variation in carriage duration [10] and propensity to cause invasive disease [11]. Most capsules are covalently attached to the peptidoglycan cell wall by Wzg (or Cps2A) enzymes [8]. The exceptions are the “mucoid” capsules 3 and 37, which are instead anchored to the membrane through attachment to phosphatidylglycerol [12]. Their mucoid appearance results from polymerases that operate continuously until synthesis is stopped by low substrate concentrations, causing the serotypes’ distinctive large colony morphologies [13].

The burden of pneumococcal disease motivated the development of polysaccharide conjugate vaccines (PCVs) that target a subset of serotypes [14]. These formulations comprise polysaccharides attached to carrier proteins, enabling infants to mount an effective adaptive immune response to the capsule antigens [15]. The increased antigenicity of protein-associated polysaccharides was first demonstrated with the serotype 3 capsule, which was attached to a horse serum globulin and used to induce protective immunity in rabbits in the 1930s [16]. Yet the first national immunisation campaign with a PCV targeting pneumococci was not initiated until 2000 [1]. This initial 7-valent design (PCV7) was later expanded as PCV10 and PCV13 formulations, which were introduced in 2010 [14] and are currently used in 148 countries [17].

PCV13 was the first to include serotype 3, which was added due to it commonly causing invasive disease associated with a high mortality rate [18]. Although PCV13 has proved highly effective in eliminating other vaccine-targeted serotypes [14], serotype 3 infections have persisted, with absolute incidences not substantially altered from the pre-PCV era [19]. This has meant serotype 3 remains a major cause of invasive disease in many countries, particularly in adults [19, 20]. The poor effectiveness against serotype 3 [21] is likely to reflect both the relatively low immunogenicity of this component in PCV13 [22] (and the 11-valent predecessor of PCV10 [23]), and the shedding of serotype 3 capsule polysaccharides from the pneumococcal membrane, which inhibits antibody-mediated killing of these bacteria [24].

Despite the absence of substantial PCV13-associated population dynamics at the serotype level, genomic epidemiology has revealed contemporaneous changes within the serotype 3 pneumococcal population. Although it is one of the most common serotypes in a genetically diverse pathogen, the serotype 3 capsule is predominantly associated with just a small number of strains [3]: of the 887 serotype 3 *S. pneumoniae* genomes in Pathogenwatch, 566 (63.8%) belong to GPSC12 (equivalent to multi-locus sequence type clonal complex 180) [25]. Pre-PCV13, the European and North American population of GPSC12 was dominated by Clade I [26–28]. Representatives of this genetically homogeneous clade typically shared an unusually stable prophage, ϕOXC141 [26] (as known as ϕSpn_OXC [29]). There was also little variation in the rest of the chromosome, as the interstrain exchange of sequence through transformation was barely detectable in these bacteria [30]. Genomic analyses of other pneumococcal strains (GPSC23, or PMEN2 [31]; and GPSC18, or PMEN9 [32]) have identified similar non-recombining lineages. In these pneumococci, the absence of diversification was attributable to stably-integrated MGEs that had disrupted competence genes required for transformation. These lineages underwent rapid local dissemination, prior to elimination by vaccine-induced immunity, as *cps* locus diversification was not observed in the absence of recombination. Analogously, in recent years, there has been an increase in non-Clade I serotype 3 GPSC12 isolates [27, 28]. However, phenotypic assays of isolates from the USA did not reveal any change in capsule expression that could directly link these changes to PCV-induced immunity [27], and the competence genes appear intact in Clade I [26, 27]. Therefore we undertook a genomic analysis of the GPSC12 population to understand how variation in recombination rates and mechanisms might have affected the epidemiological dynamics of GPSC12 in the post-PCV13 era.

## Results

### GPSC12 can be divided into distinct clades

Previous genomic analyses of serotype 3 isolates were synthesised to generate a dataset of 978 isolates, of which 971 were of sufficiently high quality to be assigned to GPSCs (Fig. S1). This identified 891 GPSC12 isolates (Fig. S2) from 24 countries (Fig. S3), with the majority (73.3%) from the UK or USA. In agreement with previous analyses of GPSC12, the recombination-corrected GPSC12 phylogeny demonstrated the strain was divided into multiple distinct clusters of isolates (Fig. 1, https://microreact.org/project/5XWnbHawQRpJyCx9aYWnnE-gpsc12-recombination-dynamics) [26–28]. In the original recombination-corrected phylogeny of GPSC12, isolates were divided into Clades I, II and III [26]. In an expanded dataset, Clade II was found to form a paraphyletic cluster subtending Clade I in a tree calculated from an alignment of core genome polymorphisms. Consequently these three clades were renamed Clades I-ɑ, I-β and II, respectively [27]. However, this reclassification appears to be contingent upon using maximum likelihood phylogenies that were not corrected for recombination [27, 28], as trees accounting for sequence exchanges did not support the renaming [26, 33]. Accordingly, neither a neighbour-joining analysis of raw genetic distances, nor a recombination-corrected maximum-likelihood phylogeny, supported Clades I-ɑ and I-β sharing a common ancestor that is exclusive of other isolates in this dataset (Fig. 1A, Fig. S2). Hence the initial classification into Clades I-III was expanded to six clades (Clades I-VI; Table S1), enabling the description of the greater diversity within GPSC12 revealed by recent genomic surveillance of the strain [28].

**Figure 1.**
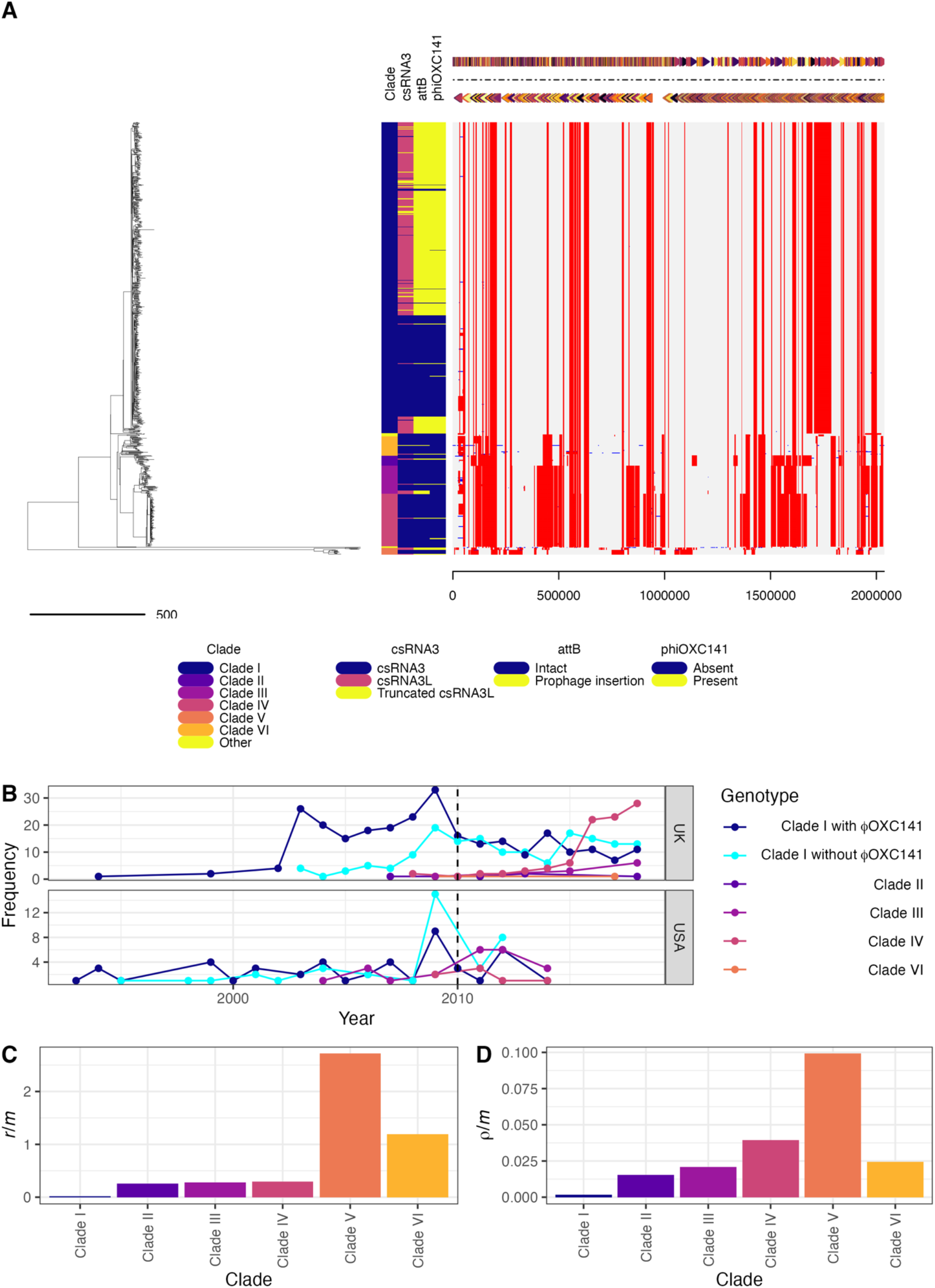
Epidemiology of the *S. pneumoniae* strain GPSC12. (A) Evolutionary reconstruction of GPSC12. The left panel shows a recombination-corrected maximum-likelihood phylogeny of GPSC12. The adjacent columns describe the distribution of genetic characteristics across the collection, with one row per isolate, according to the legend at the bottom of the panel. The leftmost column assigns isolates to clades (which can be interactively visualised at https://microreact.org/project/5XWnbHawQRpJyCx9aYWnnE-gpsc12-recombination-dynamics); the next column displays whether the unmodified csRNA3 or phage-modified csRNA3L was detected in the isolate; the next column shows whether *attB*_OXC_ is intact, or disrupted, consistent with a prophage insertion; and the rightmost column shows whether a ϕOXC141-like prophage is present in the isolate. The grey panel shows the distribution of inferred recombination events across the strain using a grid in which each row corresponds to an isolate, and each column a base in the reference genome, the annotation of which is displayed across the top. The recombination events are coloured red if they were reconstructed as occurring on an internal branch, and therefore are shared by multiple isolates through common descent, or blue if they were reconstructed on a terminal branch, and are therefore unique to a single isolate. (B) Temporal trends in the frequency of genotypes in the UK and USA, divided by assignment to clade. Clade I isolates are further subdivided by whether or not they carry a ϕOXC141-like prophage. The vertical dashed line shows the point at which PCV13 was introduced in the UK and USA. (C) and (D) Recombination dynamics within GPSC12. The extent of recombination inferred to occur within each clade is quantified as the ratio of base substitutions introduced by recombination, relative to the number occurring through point mutation (*r*/*m*), in (C), and as the number of recombination events relative to point mutations (ρ/*m*) in (D).

The majority (71.9%) of isolates belonged to Clade I (Fig. S5), which comprised representatives from every inhabited continent, and included the reference genome, *S. pneumoniae* OXC141 [26]. Phylodynamic analyses estimated this clade originated around 1954 (95% credible interval: 1938 - 1964) (Fig. S4). Most Clade I isolates retained the ϕOXC141 prophage (Fig. 1A, Fig. S6) integrated at the *attB*_OXC_ insertion site (Fig. 1A, Fig. S7) [29], despite experimental evidence of phage targeting this site also integrating at an alternative secondary location [34]. However, in addition to the known sporadic loss of the prophage [26, 27], a 210-isolate ϕOXC141-negative subclade was revealed within Clade I (32.8% of the clade; Fig. 1A). Yet there were apparent cases of ϕOXC141 reacquisition within this clade, and the presence of ϕOXC141-like prophage in other clades suggested effective within-strain transmission of this virus, as similar elements are very rare outside of GPSC12 (Fig. S8). Examples of ϕOXC141-like prophage were even identified in the distantly-related Clade V, detected in South-East Asia, containing isolates that have diverged from Clade I over centuries [33]. The other clades defined from the tree were more closely-related to Clade I. The previously-defined Clade II was split into Clades II and VI, the latter of which was commonly isolated in China and South America. Similarly, within the globally-disseminated Clade III, Clade IV was identified as mainly consisting of isolates from the UK (Fig. 1A).

### Ongoing diversification of GPSC12 in the UK and USA

The systematically-collected UK dataset [28] allowed the pre- and post-PCV13 clade prevalences to be analysed, and compared with the smaller longitudinally-collected dataset from the USA (Fig. 1B). In the UK, Clade IV rose gradually in frequency following the introduction of PCV13 in 2010, then jumped in prevalence 3.7-fold to have a frequency similar to that of Clade I in 2015. The prevalence remained elevated in subsequent years, consistent with the patterns reported previously [28].

Yet changes within Clade I were also evident in the UK pre-PCV13. From the earliest available timepoint, ϕOXC141-negative Clade I isolates increased in frequency within the UK, until they reached a similar prevalence to that of the ϕOXC141-positive Clade I isolates by the time PCV13 was introduced in 2010. These changes were mainly driven by the expansion of the ϕOXC141-negative subclade, which phylodynamic analysis suggested originated around 1980 (95% credible interval: 1974 to 1985), then expanded from the late 1990s onwards (Fig. S4). Both ϕOXC141-positive and ϕOXC141-negative Clade I isolates subsequently followed similar trends in the post-PCV13 era, with evidence of similar patterns in the USA (Fig. 1B).

The reconstruction of the strain’s diversification also suggested variation in the recombination dynamics across GPSC12. To quantify divergence through homologous recombination only, statistics were calculated excluding recombinations affecting ϕOXC141 and a putative ICE within the reference genome of OXC141 [26, 30]. By both the *r*/*m* (ratio of base substitutions introduced by homologous recombination to those resulting from point mutation) and ρ/*m* (ratio of recombination events to point mutations) measures, Clade I underwent substantially less recombination than the other GPSC12 clades (Fig. 1C, 1D, Table S2), consistent with previous analyses [27, 30]. The large number of Clade I isolates in the dataset make it unlikely the absence of detected recombinations reflects insufficient sampling of these genotypes. One parsimonious explanation for the lack of recombination in Clade I, and the rise in non-Clade I GPSC12 genotypes, is that Clade I more highly expresses the serotype 3 capsule. This could both limit DNA uptake by the competence system, and render these isolates more susceptible to PCV13-induced anticapsular antibodies.

### The type 3 capsule reduces variation through transformation

When the previously-characterised Clade I representative *S. pneumoniae* 99-4038 [26] was transformed with a rifampicin resistance marker, very few colonies were recovered across multiple experiments (Fig. 2A). To ascertain whether this inability to recombine could be ascribed to the capsule, the *cps* locus was introduced into a version of the highly-transformable unencapsulated laboratory isolate *S. pneumoniae* R6, modified to remove its phase variable *ivr* restriction-modification locus [6, 35]. This donor and recipient pair mirrored that used in the original experiments that demonstrated DNA was the transforming material [36].

**Figure 2.**
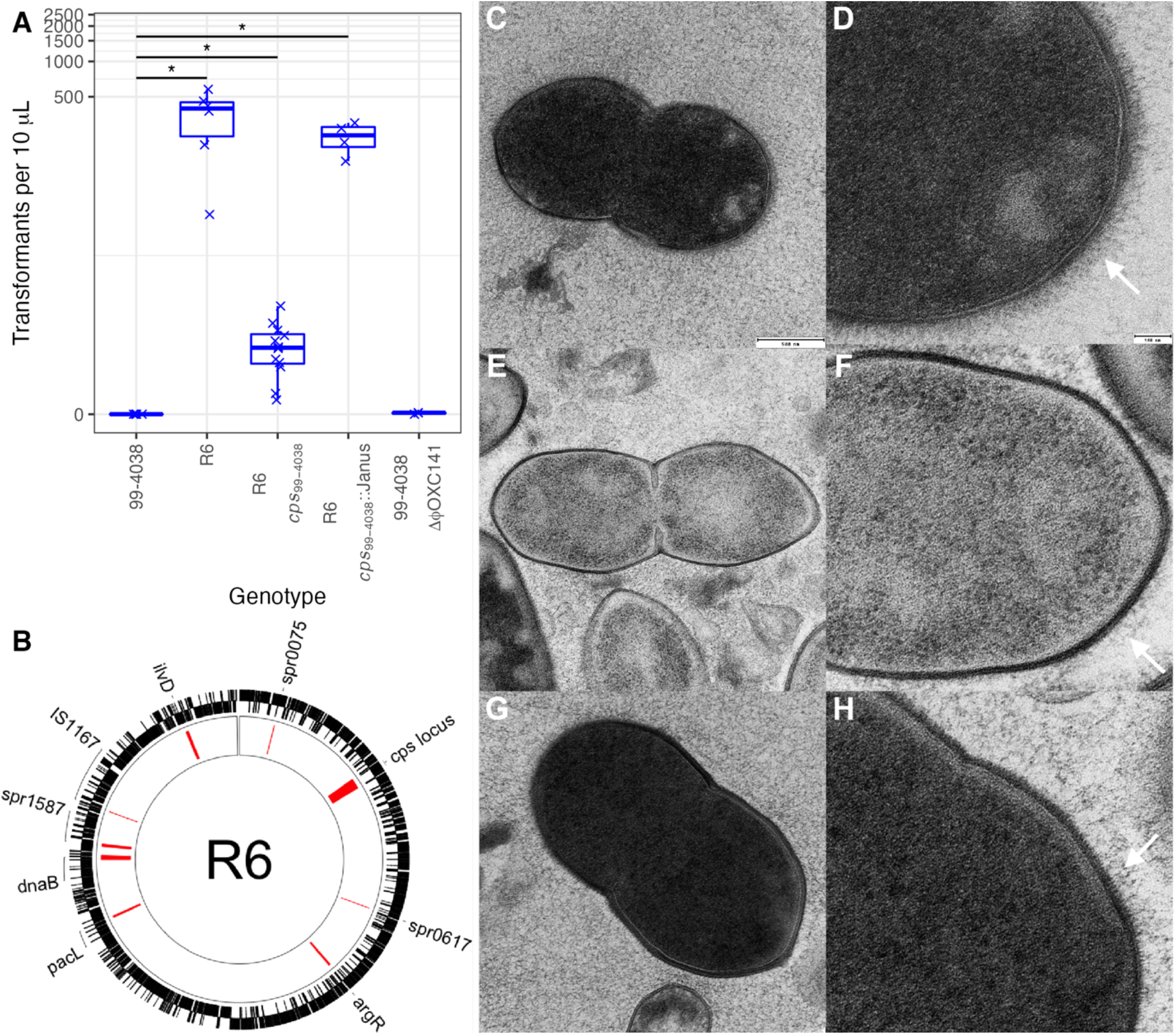
Reduction in transformation efficiency resulting from expression of the serotype 3 capsule. (A) Transformation efficiency of the Clade I isolate *S. pneumoniae* 99-4038; the unencapsulated laboratory isolate R6; the mutant R6 *cps*_99-4038_ (which carries the *cps* locus of *S. pneumoniae* 99-4038); the mutant R6 *cps*_99-4038_::Janus (in which the *cps* locus has been replaced by a resistance marker), and the mutant 99-4038 ΔϕOXC141 (in which the prophage was lost after exposure to mitomycin C). Each point represents an independent experiment, with the overall medians and interquartile ranges summarised by the box plots. A two-tailed Wilcoxon rank-sum test was used to compare the transformation frequencies of the genotypes to that of 99-4038, using a Holm-Bonferroni correction for multiple testing. Significance is coded as: *p* < 0.05, *; *p* < 0.01, **; *p* < 10^−3^, ***; *p* < 10^−4^, ****. (B) Recombinations inferred within R6 *cps*_99-4038_. The annotation of the recipient genome *S. pneumoniae* R6 is shown as a ring. The red blocks in the inner ring show the positions of inferred recombinations. Genes overlapping with these events are annotated around the edge of the panel. (C-H) Transmission electron microscopy showing the morphology of *S. pneumoniae* 99-4038, R6, and R6 *cps*_99-4038_. The panels show *S. pneumoniae* 99-4038 (C) whole cell and (D) cell surface morphology; *S. pneumoniae* R6 (E) whole cell and (F) cell surface morphology; and *S. pneumoniae* R6 *cps*_99-4038_ (G) whole cell and (H) cell surface morphology. The whole cell morphologies are shown at a consistent scale, relative to the indicated 500 nm bar. The cell surface morphologies are each shown relative to the 100 nm bar, with white arrows pointing to the layer likely representing wall teichoic acid.

One recombinant, *S. pneumoniae* R6 *cps*_99-4038_, was isolated and characterised through whole genome sequencing using the MinION platform. Analysis of the recombinant identified nine recombination events in the R6 *cps*_99-4038_ genome (Fig 2B). The largest of these was a 27,760 bp event that spanned the entire *cps* locus, importing the intact serotype 3 allele to replace the defunct serotype 2 locus in the recombination recipient (Fig. S9).

The expression of the serotype 3 capsule was demonstrated using both light (Fig. S10) and transmission electron microscopy (Fig. 2C-H). The electron micrographs revealed *S. pneumoniae* R6’s surface featured both a darkly-stained cell wall and a layer of fine threads extending tens of nanometres further out, likely corresponding to wall teichoic acid (Fig. 2F) [37]. These structures were also evident in 99-4038 and R6 *cps*_99-4038_. However, these cells were surrounded by an additional diffuse, yet extensive, matrix of capsule polysaccharides that filled the intercellular spaces (Fig. 2D, 2H). Hence R6 *cps*_99-4038_ expressed a thick capsule that replicated that of a natural serotype 3 isolate.

Transformation assays with R6, R6 *cps*_99-4038_, and a mutant in which the *cps* locus had been eliminated (*cps*_99-4038_::Janus) found expression of the capsule reduced transformation efficiency by ∼100-fold (Fig. 2A). Yet R6 *cps*_99-4038_ was still detectably transformable. Hence the type 3 capsule may contribute to the low *r*/*m* of GPSC12 overall, but is not a sufficient explanation for the absence of transformation observed in Clade I isolates *in vitro* or in epidemiological analyses. Given the precedent of mobile elements inhibiting pneumococcal transformation, the ϕOXC141 prophage was the next locus considered as a candidate for explaining Clade I’s genetic homogeneity.

### Variation in csRNA3 is commonly driven by prophage insertion at *attB*_OXC_

The *attB*_OXC_ site at which ϕOXC141 inserts was originally identified as intergenic [29], but has subsequently been shown to be within the *ccnC* gene, which generates the antisense non-coding RNA csRNA3 [34]. This is one of the *ccnA*-*E* genes in *S. pneumoniae*, encoding csRNA1-5, all of which inhibit expression of *comC* [38]. The *comC* gene encodes the protein that is processed to generate competence stimulating peptide (CSP), a quorum-sensing pheromone necessary for pneumococci to undergo efficient transformation [39]. By limiting the production of CSP, csRNAs can delay, or even block, the induction of competence for transformation [38,40,41]. Normally, insertion into the *ccnC* gene would be expected to cause a loss-of-function mutation, although the redundancy between csRNA genes would mean the phenotypic impact would likely be negligible [41]. However, the integration of ϕOXC141 instead generated two modified *ccnC*-like sequences: *ccnCL* at the *attL* site (encoding csRNA3L), and *ccnRC* at the *attR* site (encoding csRNA3R; Fig. S11), as observed for phage ϕSpSL1 [34]. The *ccnCL* gene is expected to be functional, as its promoter is that of the native *ccnC* gene, and the 5’ stem-loop of csRNA3L is unmodified. However, the 3’ stem-loop is changed from having four unpaired bases in the loop, to instead comprising the sequence CAAUCA (Fig. S12). This matches the sequences of the 3’ loops in csRNA1, csRNA3 and csRNA4, each of which is more effective at inhibiting competence than the unmodified csRNA3 allele [40]. This does not changes the predicted affinity of the csRNA3 for the *comC* transcript (Fig. S12), but may alter its interaction with Cbf1, which seems to stabilise and process these RNAs [42]. Hence ϕOXC141 insertion has the potential to increase the effectiveness of a cell’s csRNA repertoire at preventing competence induction, thereby accounting for the reduced rate of homologous recombination observed in Clade I (Fig. S13).

To ascertain whether the modification of csRNA was common across the species, the 20,047 pneumococcal genomes from the GPS project were scanned for *ccn* genes [3]. This identified 104,934 csRNA sequences, with a mode of five per genome (Fig. S14). A phylogeny of a non-redundant set of 388 sequences enabled the majority of csRNA alleles to be classified into eight sets (Fig. 3A; Table S3). Three of the csRNAs (csRNA2, csRNA4 and csRNA5) were present in >98% of isolates, exhibiting very little variation across the species (Fig. 3B). The csRNA1-type sequences were only present in ∼85% of isolates. The corresponding *ccnA* gene is found in tandem with *ccnB* (encoding csRNA2) upstream of *ruvB* [40]. An independent analysis of this chromosomal locus found it was structurally divergent from the version in *S. pneumoniae* R6 in ∼18% of isolates (Fig. S15). Within Clade I, a comparison of the complete genomes of *S. pneumoniae* OXC141 and 99-4038 suggested this was the result of an intragenomic recombination between *ccnA* and *ccnB* (Fig. S15). The chimeric *ccnAB* sequences produced by these events were classified as csRNA2, as they retained the 5’ stem-loop of csRNA1, which is similar between both genes, and the 3’ stem-loop of csRNA2, which is divergent from that of csRNA1 (Fig. 3B).

**Figure 3.**
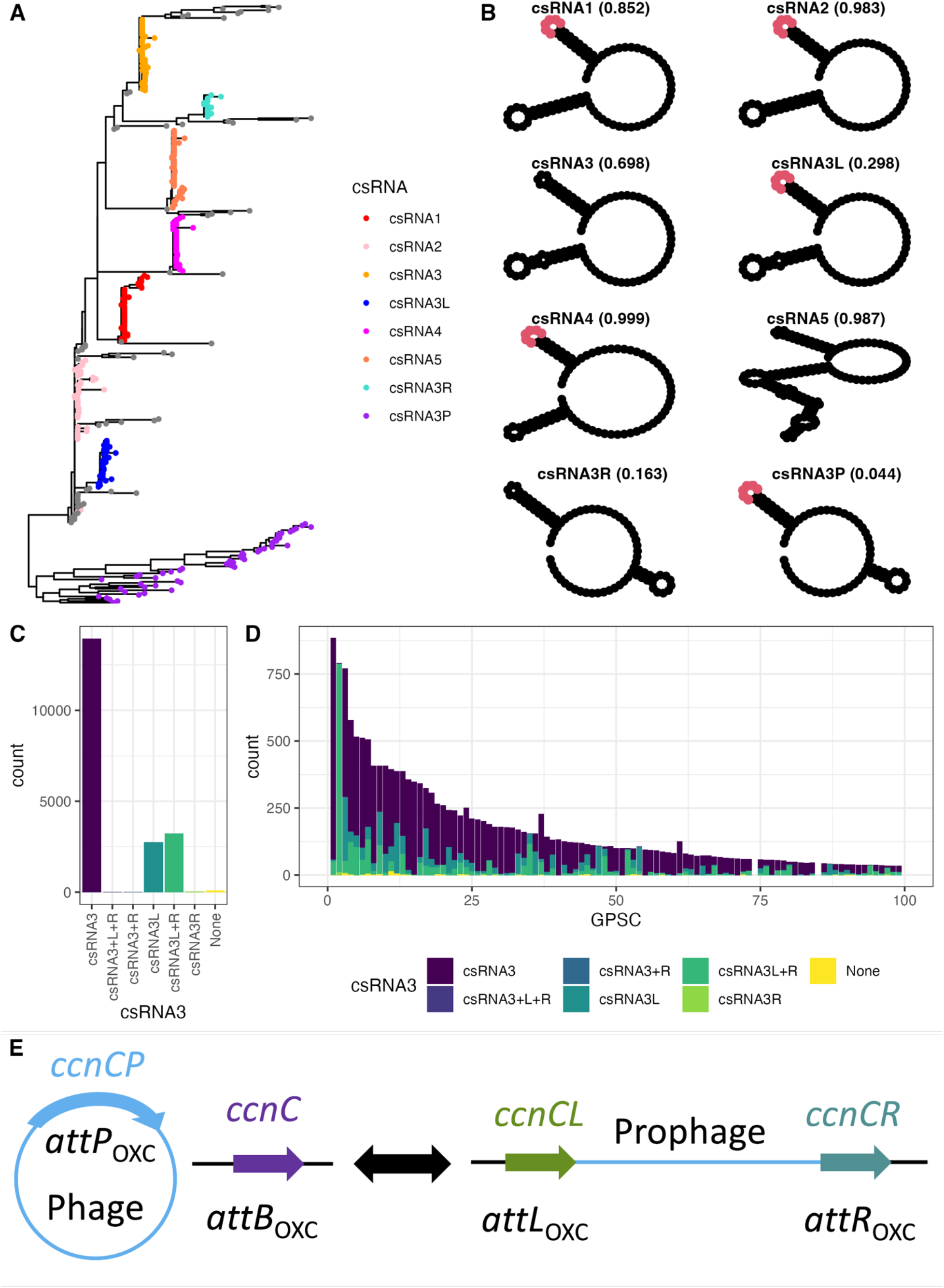
Distribution of csRNA sequences across the 20,047 Global Pneumococcal Sequencing (GPS) project isolates. (A) A maximum-likelihood phylogeny of 388 csRNA sequences, representing the non-redundant set identified within the GPS collection after excluding examples <80 nt in length. Nodes are coloured according to assignment to the eight different csRNA types described in the text. (B) The predicted structure of the most common sequence of each csRNA type is shown. The sequence CAAUCA is highlighted in red where it is present. The total frequency of all sequences assigned to the named allele in the GPS isolates is shown in parentheses. (C) The distribution of csRNA3 allele combinations across the GPS collection. The bar chart quantifies the prevalences of different combinations of the unmodified csRNA3 type, and the prophage-modified csRNA3L and csRNA3R types, across isolates. (D) Distribution of different csRNA3 type combinations across Global Pneumococcal Sequence Clusters 1 to 100 in the GPS collection. (E) Model for the interconversion between csRNA3P (encoded by *ccnCP* at *attP*) and csRNA3 (encoded by *ccnC* at *attB*), when a phage is excised from the chromosome, and csRNA3L (encoded by *ccnCL* at *attL*) and csRNA3R (encoded by *ccnCR* at *attR*), when the prophage is integrated.

The csRNA3 and csRNA3L sequences were present in 69.8% and 29.8% of isolates respectively, but never found within the same isolate (Fig. 3C). This indicates almost all isolates had one of these two mutually-exclusive alleles. Over half of isolates with a csRNA3L were also found to have a csRNA3R (present in 16.3% of isolates), which was almost never found in the absence of csRNA3L. The inability to detect csRNA3R in many csRNA3L-positive genomes may represent the reduced functional constraints on csRNA3R sequences, as their predicted structure features a short stem-loop that suggests they may not form an effective csRNA (Fig. 3B). The same small stem-loop is present on a representative of a diverse set of sequences labelled csRNA3P, which are most divergent from the previously-characterised csRNA sequences (Fig. 3B). The corresponding *ccnCP* genes can be found at the *attP* sites of phage genomes, between the integrase and lytic amidase genes (Fig. S16) [34]. Correspondingly, the csRNA3P stem-loop that does not match with csRNA3R is that which replaces the 3’ stem-loop of csRNA3 to generate csRNA3L (Fig. 3B). Hence divergent *ccnCP* sequences carried on diverse prophage modify the cellular *ccnC* gene through insertion to generate *ccnCL* and *ccnCR* genes (Fig. 3E). The widespread alteration affects approximately one-third of pneumococci.

### Modification of csRNA3 in serotype 1 and genetically-tractable genotypes

To test whether the integration of ϕOXC141 inhibited the transformation of Clade I, *S. pneumoniae* 99-4038 was passaged in the presence of the lysogen inducer mitomycin C [43], enabling the isolation of bacteria lacking the prophage (ΔϕOXC141; Fig. S17). When competence was induced by exogenous CSP (Fig. S13), the transformation efficiency of ΔϕOXC141 was similar to that of the parental genotype (Fig. 2A). This suggested 99-4038 would be a poor model for understanding the effects of csRNA3 alteration, as this change was hypothesised to inhibit endogenous CSP production, the quantification of which requires lower-efficiency assays of spontaneous transformation. Nevertheless, the frequency of the *ccnCL* in the pneumococcal population made it possible to search for a genetically tractable isolate carrying this prophage-modified sequence.

The *ccnCL*-type sequences were detected in almost half of GPSCs (292 of 586; 49.8%), and were almost ubiquitous within GPSC2, the most common strain expressing serotype 1 (Fig. 3D) [3]. This strain is responsible for a high proportion of invasive disease in Africa, and is regarded as being unusually genetically homogenous [3,44,45]. The modification of csRNA3 results from the insertion of ϕPNI0373 at *attB*_OXC_ (Fig. S18) [46]. Analysis of the GPS collection suggests this prophage is conserved between almost all GPSC2 isolates, despite being absent from all other GPSCs (Fig. S19). Yet generating mutant variants of serotype 1 pneumococci is challenging [46]. Instead, the effects of csRNA3L were investigated in GPSC97 isolate *S. pneumoniae* RMV8, which can be efficiently transformed [47, 48]. Although this isolate has a CSP2 pherotype, rather than the CSP1 pherotype of Clade I, the interactions of both csRNA3 and csRNA3L were predicted to be similar with the *comC* transcripts generating both CSP types (Fig. S12).

The modification of *ccnC* in *S. pneumoniae* RMV8 resulted from the integration of a 32.8 kb prophage, ϕRMV8, at *attP*_OXC_ (Fig. S20). Previous experimental work on this genotype resulted in the isolation of two variants, each expressing different alleles of the phase-variable SpnIV restriction-modification system [47], encoded by the translocating variable restriction (*tvr*) locus [4]. Further alterations at the *tvr* loci of these isolates were precluded by the disruption of the *tvrR* gene, encoding the recombinase responsible for rearrangements through excision and integration (Fig. 21) [47]. The more common “dominant” variant (RMV8_domi_) expressed a SpnIV allele that methylated the motif GATAN_6_RTC, whereas the allele expressed by the less common “rare” variant (RMV8_rare_) methylated the motif GTAYN_6_TGA. Comparing the growth curves of these variants, and *tvr*::*cat* derivatives in which the *tvr* locus was replaced with a chloramphenicol resistance marker, had found RMV8_rare_ exhibited a growth defect in late exponential phase (Fig. S21). This correlated with increased activity of ϕRMV8 in RMV8_rare_ during stationary phase, and correspondingly the growth defect was not observed in an RMV8_rare_ ϕRMV8::*tetM* mutant (Fig. S21). Hence the distinctive growth profile of RMV8_rare_ was the consequence of increased activation of ϕRMV8. This correlated with different epigenetic modification of the pneumococcal chromosome, analogous to the variation in PRCI activity seen between *tvr* variants in another pneumococcal genotype [48].

### Prophage-driven alteration of csRNA3 affects *comC* expression

To understand how differences in ϕRMV8 activity might affect csRNA3-type sequences, RNA-seq data were generated for three replicates of each of the four RMV8 genotypes (the two variants, and the corresponding *tvr*::*cat* mutants) during the late exponential growth phase (Table S4). The sequence reads exhibited consistent inferred fragment size (Fig. S23) and gene expression (Fig. S24) distributions, suggesting the datasets should be informatively comparable. Comparing RMV8_domi_ with the other three genotypes using Q-Q (Fig. S25) and volcano (Fig. S26) plots suggested the main differences between the genotypes could be captured with a false discovery rate threshold of 10^-3^ (Table S5). As a positive control, this found the *tvr* loci to be more highly expressed in RMV8_domi_ than either of the genotypes in which these genes were deleted (Fig. 4). The only two other loci to exhibit significant differences in transcription were ϕRMV8 and *comC* (Table S6). Consistent with the growth inhibition attributable to elevated ϕRMV8 activity in RMV8_rare_, the prophage was more highly transcribed in this genotype across the lysogeny (Fig. S27), replication (Fig. S28), structural (Fig. S29) and lysis (Fig. S30) genes, although not all reached statistical significance (Fig. 4A). This heightened prophage activity was not evident in RMV8_rare_ *tvr*::*cat*, which instead upregulated *comC* relative to the other genotypes (Fig. 4A). The co-operonic *comDE* genes, encoding the CSP receptor, were also more highly transcribed in this genotype (Fig. S31). Hence the RNA-seq data demonstrated a specific link between a prophage at *attB*_OXC_ and transcription of the *comCDE* operon.

**Figure 4.**
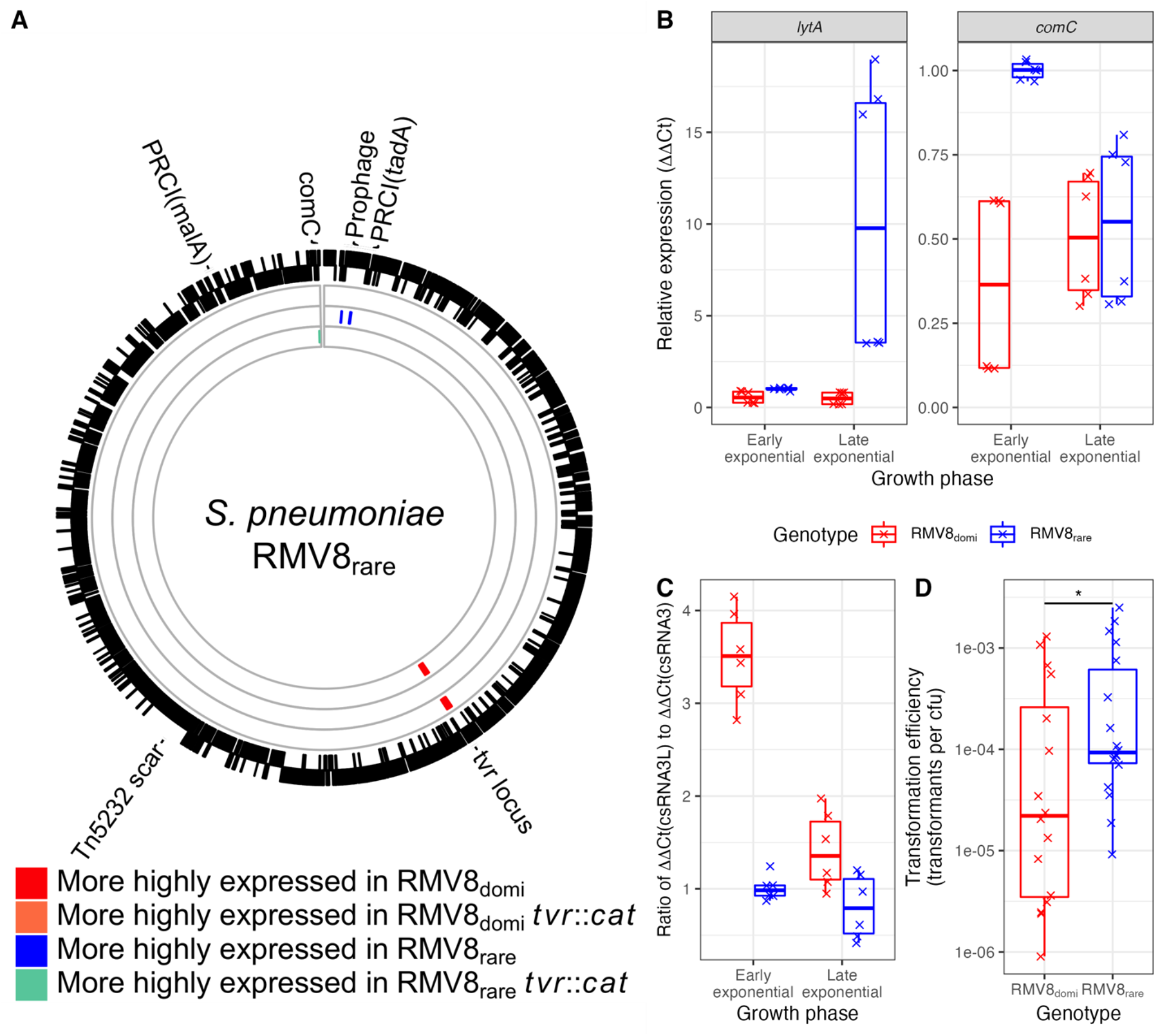
Variation between RMV8 genotypes. (A) RNA-seq analysis of *S. pneumoniae* RMV8. The outer rings represent the annotation of the complete genome of the *S. pneumoniae* RMV8_rare_ variant (accession code OX244288). The mobile elements present in this isolate, corresponding to a prophage, two phage-related chromosomal isolates (PRCIs; inserted near *tadA* and *malA* [4]) and a Tn*5253* scar [98] are marked. The inner rings show the differences in expression between RMV8_domi_, the most frequently-isolated genotype in RMV8 cultures, and derived genotypes. The outermost of these rings shows the comparison with RMV8_domi_ *tvr*::*cat*, in which the *tvr* locus was replaced with a *cat* resistance marker. The next ring shows the comparison with RMV8_rare_, in which significant differences at the prophage locus were detected. The innermost ring shows the comparison with RMV8_rare_ *tvr*::*cat*, which highly expressed the *comC* gene. (B) Quantification of ϕRMV8 *lytA* and *comC* expression by qRT-PCR in RMV8_domi_ and RMV8_rare_ samples collected at early (OD_600_ = 0.2) and late (OD_600_ = 0.5) exponential phase growth. Six points are shown, corresponding to three technical replicates measurements of two biological replicates. Data are coloured according to the genotype being tested. (C) Quantification of the relative levels of csRNA3L and csRNA3 in early and late exponential phase growth. Data are shown as in panel (B). (D) Scatterplot showing the spontaneous transformation efficiency of the RMV8 variants. Each point represents an independent experiment, with the overall medians and interquartile ranges summarised by the box plots. A two-tailed Wilcoxon rank-sum test was used to compare the transformation frequencies of the variants, using a Holm-Bonferroni correction for multiple testing. Significance is coded as: *p* < 0.05, *; *p* < 0.01, **; *p* < 10^−3^, ***; *p* < 10^−4^, ****.

As conventional RNA-seq library preparation does not efficiently incorporate small non-coding RNAs, understanding whether csRNA3 linked the changes in *comCDE* and ϕRMV8 transcription required quantitative reverse-transcriptase PCR (qRT-PCR) experiments. Samples were taken during early exponential growth (OD_600_ = 0.2), when competence is typically inducible, and late exponential growth (OD_500_ = 0.5), when the RNA-seq samples were collected. Comparisons of RMV8_domi_ and RMV8_rare_ confirmed the ϕRMV8 *lytA* gene was more active in the latter genotype (Fig. 3B). The increased excision of ϕRMV8 in RMV8_rare_ (Fig. S21) raised the expression of csRNA3, from the restored *ccnC* gene, relative to csRNA3L (Fig. 3C). This ratio was highest during early exponential phase, and correspondingly *comC* expression in RMV8_rare_ was approximately twice that of RMV8_domi_ at this growth stage (Fig. 3B). This was sufficient to significantly alter the induction of competence, as RMV8_rare_ underwent spontaneous transformation four-fold more frequently than RMV8_domi_ (Fig. 4D), despite there being no significant difference in their transformability when CSP was added exogenously [48]. Hence more stably-integrated prophage correlated with a higher ratio of csRNA3L to csRNA3 expression, lower *comC* transcription, and reduced transformability.

Consistent with the RNA-seq data, *comC* expression was similar in RMV8_domi_ and RMV8_rare_ in late exponential phase (Fig. 3B), but elevated in RMV8_rare_ *tvr*::*cat* (Fig. 32). This appeared to reflect the low levels of csRNA3 and csRNA3L in RMV8_rare_ *tvr*::*cat* during early exponential phase (Fig. S32), which correlated with the low activity of ϕRMV8 in this genotype (Fig. S21 and S32). However, the measurement of csRNA3L levels and ϕRMV8 transcription by qRT-PCR were not independent. Mapping RNA-seq data to the *attL*_OXC_ site demonstrated antisense expression of *ccnCL* was driven by transcription initiated within ϕRMV8 while it was integrated in the chromosome (Fig. S32). This has the potential to confound the quantification of csRNA3L levels in lysogenic genotypes, as the variable intracellular dynamics of prophage excision and reintegration changed the expression of csRNA3 and csRNA3L between even near-isogenic cells.

### Expression of csRNA3L inhibits activation of transformation through quorum sensing

To establish whether csRNA3L caused the differences in transformation efficiency required the construction of mutants stably expressing one allele at the *attB*_OXC_ site. Both *ccnCL* and ϕRMV8 were replaced with a single fixed csRNA3-type sequence in RMV8_rare_ and RMV8_domi_: either *ccnC* from R6 or *ccnCL* from RMV8_rare_. No substantial growth difference was observed between the two genotypes *in vitro*. Spontaneous transformation assays demonstrated both *ccnC* mutants exhibited significantly higher transformation efficiencies than the corresponding *ccnCL* mutants: an 11-fold difference in RMV8_rare_, and an eight-fold difference in RMV8_domi_ (Fig. 5A). Negative control experiments, conducted in the absence of DNA or the presence of DNase I, confirmed these assays should accurately reflect levels of spontaneous transformation (Fig. S33)

**Figure 5.**
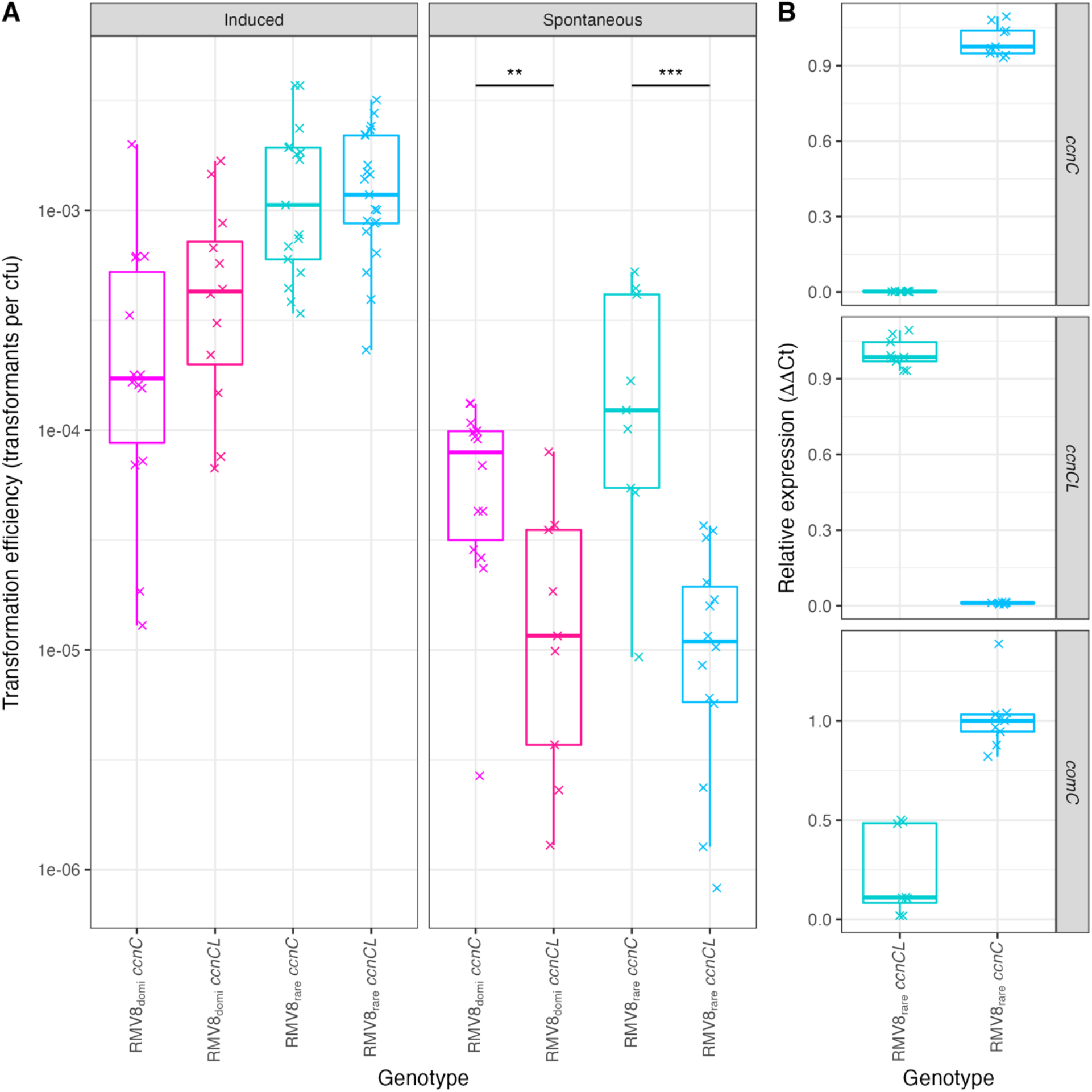
The increased inhibition of competence induction by *ccnCL* relative to *ccnC* in RMV8. (A) Transformation frequencies of independent RMV8_rare_ and RMV8_domi_ mutants expressing *ccnC* and *ccnCL*. Frequencies were calculated as the count of rifampicin-resistant colonies relative to the number of cells in the sample. Each point represents an independent experiment, summarised for each genotype as a boxplot. Data are coloured according to genotype. Induced competence experiments measured the frequency of recombination following the addition of exogenous CSP. Spontaneous competence experiments measured the frequency of recombination in overnight cultures with no added CSP. A two-tailed Wilcoxon rank-sum test was used to compare the transformation frequencies of the matched *ccnC* and *ccnCL* mutants within each background, using a Holm-Bonferroni correction for multiple testing. Significance is coded as: *p* < 0.05, *; *p* < 0.01, **; *p* < 10^−3^, ***; *p* < 10^−4^, ****. (B) Levels of *ccnC*, *ccnCL* and *comC* transcripts in RMV8_rare_ mutants expressing either *ccnC* or *ccnCL* at the end of spontaneous transformation experiments. RNA concentrations were estimated using qRT-PCR, and quantified using the ΔΔCt approach (see Methods). The points represent the data from three technical replicates measurements of each of three biological replicates.

By contrast, when competence was induced with exogenous CSP during early exponential phase growth, no significant difference was observed between the mutants in the RMV8_rare_ or RMV8_domi_ backgrounds (Fig. 5A). Hence the observed differences between the *ccnC* and *ccnCL* genotypes did not represent variation in the activity of the competence machinery itself. To test the hypothesised causal mechanism, the expression of csRNA3, csRNA3L and *comC* was quantified in the RMV8_rare_ mutants under the conditions used to assay spontaneous transformation (Fig. 5B). These confirmed csRNA3 and csRNA3L were only detectable in the corresponding mutants. They also demonstrated that *comC* expression was reduced more than four-fold in the *ccnCL* mutant relative to the *ccnC* mutant. Therefore prophage-mediated RNA alteration inhibited the ability of pneumococci to spontaneously induce competence through the endogenous production of a quorum sensing signal.

## Discussion

The pneumococcus has proved adept at acquiring antibiotic resistance and evading vaccine-induced immunity through transformation [49]. Yet GPSC12 isolates have persisted as a major cause of disease after the introduction of PCV13 while retaining their serotype 3 capsule and typically remaining pansusceptible to antibiotics [3]. This absence of adaptive evolution to public health interventions is the consequence of the strain undergoing relatively little diversification through homologous recombination, with the overall *r*/*m* of 0.56 more than an order of magnitude lower than that calculated from samples of commonly antibiotic resistant strains such as GPSC1 [50], GPSC6 [32] or GPSC16 [30]. The mucoid serotype 3 capsule clearly contributes to the low recombination rate, likely through acting as a physical barrier to the acquisition of exogenous DNA. Yet this is insufficient to explain the low frequency of recombinations in Clade I relative to the rest of GPSC12.

Clades with a greater propensity to acquire diverse sequence through recombination than Clade I have been increasing in frequency in the UK and USA post-PCV13 [27, 28]. There is no evidence that this can be directly ascribed to differential effects of vaccine-induced immunity, based on physiochemical and serological assays applied to different clades previously [27]. Yet given that Clade I is decades old [26, 27], and the composition of GPSC12 changed rapidly in the UK just as the prevalence of other strains was shifting following the introduction of PCV13 [51], it is likely that these changes reflect the indirect consequences of vaccine-induced immunity on the bacterial population [52].

However, there has also been a change within Clade I itself that pre-date PCV13’s introduction. This has been driven by the expansion of a subclade lacking ϕOXC141, suggesting this element is deleterious to its host cell, albeit not causing such a large cost that would drive a rapid selective sweep of ϕOXC141-negative genotypes. The monophyletic expansion of the large ϕOXC141-negative clade suggests the prophage is infrequently lost [53], consistent with the rarity of ϕOXC141-negative isolates in mitomycin C passages (Fig. S17). The ϕNPI0373 prophage appears to be similarly stably integrated in the *attB*_OXC_ site of GPSC2. GPSC2 and GPSC12 both also show little evidence of recent diversification through transformation [45]. Hence these “frozen” genotypes contrast with other pneumococcal strains that undergo more frequent recombination, in which individual prophage infections are typically ephemeral [4, 7].

In GPSC2, the lack of transformation has been partially attributed to disruptive mutations in competence genes [46]. Similarly, the lack of diversification through homologous recombination in Clade I cannot be entirely explained by the phage-mediated alteration of *ccnC*. While the serotype 3 capsule substantially limits the efficiency of DNA import, transformants were still detectable in an R6 mutant expressing this capsule following addition of exogenous CSP, in contrast to *S. pneumoniae* 99-4038. This implies there is a difference in transformation efficiency that is independent of both the capsule and the ability to generate endogenous CSP, reflecting the complexity of the regulation of transformation [5, 48]. The relative importance of these multiple inhibitory mechanisms may be quantified by the ϕOXC141-negative subclade within Clade I, which has a restored *ccnC* (Fig. 1A). There is little evidence yet of an increased transformation rate in this group, but this is unsurprising, given its recent expansion, limited geographic sampling, and the low *r*/*m* across GPSC12. Yet if its dissemination continues, the rate of recombination compared to the rest of Clade I might reveal the importance of *ccnCL* relative to other aspects of the GPSC12 genotype.

Nevertheless, the alteration of csRNA3 is likely to be hugely influential on pneumococcal evolution at a broader scale, owing to this polymorphism being found in almost half of strains, and almost one-third of individual pneumococcal isolates. Perhaps the most persuasive evidence of its importance is the conservation of the phage sequences contributing to the *ccnCL* allele, in contrast with the otherwise immensely variable nature of pneumococcal prophage [4,29,55]. This is indicative of a strong selective benefit of the csRNA3 alteration to the virus. Through limiting transformation within communities of cells, prophage generating *ccnCL* alleles can limit the rate at which they are deleted as they spread between previously-uninfected cell populations within a host nasopharynx [7, 54]. Notably, the *ccnCL* allele is only present when the prophage is integrated into the chromosome, and therefore vulnerable to deletion through chromosomal curing, with the original *ccnC* restored on phage excision (Fig. 3E). Hence the alteration of the cell’s csRNA repertoire can be explained as a defence mechanism of a selfish MGE.

This is the second of the four commonly-targeted pneumococcal *attB* sites [56] at which prophage integration has been found to inhibit transformation. Although prophage insertion increased the effectiveness of the competence-inhibiting *ccnC*, similar elements cause loss of function when they disrupt the pilus gene *comYC*, which is necessary for competence [31, 49]. The truncation of *comYC* was recently demonstrated to be the most common single type of protein-modifying mutation affecting coding sequences during within-host pneumococcal evolution [57]. This emphasises the importance of conflict between prophage and the host cell in rapidly generating common polymorphisms in this bacterium [54].

While the disruption of the competence machinery by mobile elements has already been shown to be widespread across bacterial pathogens [7], the potential for blocking transformation through modification of regulatory non-coding RNAs is becoming increasingly apparent. In addition to csRNAs, another pneumococcal non-coding RNA has been found to inhibit transformation by repressing *comD* expression [58]. In *Legionella pneumophila*, the RocR non-coding RNA was identified as a repressor of competence [59]. Subsequently, a set of *L. pneumophila* mobile elements were found to carry a modified RocRp RNA, enabling the elements to inhibit transformation through a mechanism analogous to the modification of csRNA3 [60]. Hence the manipulation of competence regulatory systems as part of the conflict between horizontal DNA transfer mechanisms is likely to emerge as an important factor in shaping genetic diversity across more species as our understanding of mobile elements and transformation regulation improves. These trends must then be disentangled from the effects of vaccine and antibiotic selection to understand the epidemiology of bacterial pathogens.

## Methods

### Phylogenetic analysis of GPSC12

For the analysis of GPSC12’s population structure, an overlapping set of 1116 short-read datasets was collated from from Azarian *et al* (295 isolates mainly from Europe and the USA) [27], Groves *et al* (616 isolates collected by Public Health England) [28], and the Global Pneumococcal Sequencing (GPS) project (205 isolates from https://microreact.org/project/gpsGPSC12) [33]. These datasets yielded a non-redundant set of 979 isolates, one of which (accession code ERR433970) was inaccessible. The short read data for the remaining 978 isolates were assembled *de novo* with SPAdes version 3.10.1 [61] using default settings and k-mer lengths between 21 and 85 with a step size of four. Assemblies were evaluated with assembly-stats [62]. Seven sequences with anomalous assembly lengths, below 1.9 Mb or above 2.5 Mb, were removed.

The 971 remaining isolates assembled from short-read data, and six high-quality draft genomes from Croucher *et al* [26], were assigned to strains using PopPUNK version 2.4.0 [2] and version 6 of the Global Pneumococcal Sequencing project’s Global Pneumococcal Sequence Cluster database [3]. This found 891 of the isolates belonged to strain GPSC12, and therefore were sufficiently closely related for phylogenetic analysis.

The complete genome of GPSC12 isolate *S. pneumoniae* OXC141 (accession code FQ312027) was used as the reference genome against which all other 890 GPSC12 assemblies were mapped using SKA version 1.0 with default settings [63]. The resulting alignment of 891 whole genomes was analysed with Gubbins version 3.2.1 [64], using RapidNJ [65] and a Jukes-Cantor model to generate the original tree, with four subsequent iterations using RAxML [66] trees constructed with a General Time-Reversible model of base substitution using a gamma model of between-site rate heterogeneity. Results were visualised with Phandango [33] and RCandy [67].

### Analysis of csRNA

Identification of csRNA used the corresponding Rfam model (accession code RF02379) [68]. Assembled genomes were scanned using the cmsearch tool from version 1.1.2 of the Infernal package [69]. Heuristic filters were set at the levels used for Rfam, and both significant and questionable hits were retained, to explore the full diversity of csRNA sequences.

To analyse the diversity of csRNA sequences in the GPS dataset, the 104,934 csRNA sequences were processed to identify a non-redundant set of 388 sequences that were all longer than 80 nt and contained no ambiguous bases. These were aligned with mafft v7.505 [70] using the default progressive alignment FFT-NS-2 method with a DNA200 model. A phylogeny was generated using FastTree version 2.1.11 [71] with a Jukes-Cantor model and a CAT approximation of between-site rate heterogeneity with 20 rate categories. The phylogeny was manually annotated with FigTree [72] and ggtree [73].

RNA structures were visualised using the minimum free energy prediction [74] generated by the RNAfold WebServer [75] with default settings. Structures were plotted using the R package RRNA [76]. Inference of the strength of interactions between csRNAs and the *comC* mRNA, using a previously-identified *comC* transcription start site [77], used IntaRNA version 2.0 [78].

### Analysis of the accessory genome

The 871 assembled GPSC12 isolates were used to generate a sequence database with BLAST version 2.12.0 [79]. To identify intact *attB*_OXC_ sequences, a 4,477 bp locus was extracted from the *S. pneumoniae* TIGR4 genome (accession code AE005672) [80], extending from the 5’ end of *purA* to the 3’ end of *radA*. GPSC12 isolates with a BLASTN match longer than 3,750 bp were inferred to have an intact *attB*_OXC_ site, whereas isolates with only shorter alignments to the query sequence were inferred to have a prophage insertion at this site (Fig. S7). Similarly, the presence of prophage ϕOXC141 was inferred through querying the database with the 34,080 bp ϕOXC141 sequence from *S. pneumoniae* OXC141 [26]. If the longest BLASTN hit was longer than 12,500 bp, then an isolate was inferred to be infected with ϕOXC141, or a very closely-related prophage (Fig. S6).

Phylodynamic analysis of Clade I applied BactDating v1.1.1 [81] to the subset of 415 isolates with a date of isolation, which ranged from 2002 to 2018. The analysis used a relaxed clock model, fitted using Markov chain Monte Carlo (MCMC) sampling run for 5×10^7^ iterations. Half of the chain was discarded as burn-in, with convergence of the second half of the chain established through visual assessment of the MCMCs (Fig. S4). To estimate the date at which the ϕOXC141-negative subclade emerged, the presence and absence of ϕOXC141 was reconstructed over the internal nodes of the Gubbins phylogeny as a discrete state using PastML with the JOINT model [82].

To explain the relatively low frequency of csRNA1 sequences in the GPS collection, the 20,047 sequences described by Gladstone *et al* were used to generate a species-wide sequence database [3]. This was queried with the intergenic sequence upstream of *ruvB* in *S. pneumoniae* R6 (accession code AE007317) [35], the locus within which *ccnA* and *ccnB* were originally identified [38]. Assemblies with a BLAST alignment to the query that was longer than 900 bp were inferred to have a full-length locus encoding both *ccnA* and *ccnB*, whereas assemblies with an alignment below this threshold were assumed to have the chimeric gene *ccnAB* (Fig. S15).

Pairwise alignments of genomes for analysis of specific accessory loci used BLASTN, and were visualised using the Artemis Comparison Tool [83] and the R package genoPlotR [84].

### Culturing and passage of *S. pneumoniae*

*S. pneumoniae* were grown statically at 35 °C in 5% CO_2_, unless otherwise stated. Genotypes used in this study are listed in Table S7. Liquid cultures were grown in a mixture of 2:3 ratio of Todd-Hewitt media with 0.5% yeast extract (Sigma-Aldrich), and Brain-Heart Infusion media (Sigma-Aldrich), dissolved in milliQ (18MΩ) water, henceforth referred to as mixed media [48]. Culturing on solid media used Todd-Hewitt media with 0.5% yeast extract.

For each passage of *S. pneumoniae* 99-4038, an overnight culture was diluted 1:9 in fresh mixed media supplemented with 40 µg mL^-1^ catalase to a total volume of 10 mL. At an OD_600_ of between 0.3 and 0.4, mitomycin C was added to a final concentration of 0.1 µg mL^-1^. After three such passages, 24 colonies were picked and screened for the loss of ϕOXC141 by PCR amplification of the *attL*_OXC_ and *attB*_OXC_ sequences.

### Extraction of DNA and PCR amplification

Cultures were centrifuged for 10 min at 4,000 *g* and the supernatant was discarded. Bacterial cell pellets were resuspended in 480 μL lysis buffer (Promega) and 120 μL 30 mg mL^-1^ lysozyme (Promega), followed by 30 min incubation at 35 °C. Samples were centrifuged at 8,000 *g* for 2 min, and DNA extracted from the pellets using the Wizard Genomic DNA Purification Kit (Promega) according to the manufacturer’s instructions.

Extraction of genomic DNA from *S. pneumoniae* R6 *cps*_99-4038_ for sequencing with Oxford Nanopore technology used a different approach, to reduce the polysaccharide contamination of the final sample used for library preparation. A cell pellet was resuspended in 250 μL Tris-EDTA buffer and 50 μL 30 g L^-1^ lysozyme (Roche) in Tris-EDTA buffer. This mixture was vortexed at room temperature for 15 mins and 400 μL 0.1 M EDTA (Gibco) and 250 μL 10% sarkosyl (BDH) were added. Samples were incubated at 4 °C for 2 h, prior to the addition of 50 μL proteinase K (Roche), 30 μl RNase A (Roche) and 3 mL Tris-EDTA buffer. Samples were incubated at 50 °C overnight, then washed with 5 mL of a 25:24:1 mixture of phenol, chloroform and indole-3-acetic acid (Fluka) and centrifuged (2,594 g, 10 min). The aqueous phase was removed, washed with 5 mL chloroform (Sigma-Aldrich) and centrifuged (2,594 g, 10 min). DNA was precipitated from the aqueous phase using 300% by volume ethanol and 10% by volume 3 M sodium acetate followed by 1 h incubation at −20 ℃. The pellet recovered following centrifugation was then washed with 5 mL 70% ethanol and resuspended in water.

PCR amplification used 500 ng of template DNA, 7.5 µL 2x DreamTaq Master Mix (ThermoFisher), nuclease-free water (ThermoFisher) and 1 µl of a 10 µM solution of each of the forward and reverse primers. Primer sequences are listed in Table S8.

To purify individual DNA amplicons, PCR products were separated using 1% agarose gels (Sigma-Aldrich) dyed with SYBR Safe (Invitrogen) in TBE buffer (Invitrogen) with a 1 kb HyperLadder (Bioline) marker. Where necessary, individual amplicons were excised and extracted with the GenElute Gel Extraction Kit (Sigma-Aldrich) according to manufacturer’s instructions.

### Extraction and processing of RNA samples

To generate samples for RNA-seq, the four genotypes being analysed (*S. pneumoniae* RMV8_domi_, RMV8_domi_ *tvr*::*cat*, RMV8_rare_ and RMV8_rare_ *tvr*::*cat*) were grown as 500 µL cultures in a 48-well plate to an OD_600_ of 0.5. Four wells were then harvested at the appropriate growth stage and combined with 4 mL RNAprotect (Qiagen) to generate each replicate. These samples were then processed with the SV Total RNA Isolation System (Promega) according to the manufacturer’s instructions. The generation of cDNA and Illumina sequencing libraries was undertaken as described previously [48]. Three replicates for each genotype were sequenced as 200 nt paired-end multiplexed libraries on a single Illumina HiSeq 4000 lane.

To generate samples for qRT-PCR, cells were grown to the specified optical density in 10 mL mixed liquid media with 500 mM calcium chloride. A 5 mL sample was then mixed with 10 mL of RNAprotect and processed with the SV Total RNA Isolation System (Promega) according to the manufacturer’s instructions. DNA was removed from 0.5 μg samples of RNA with Amplification-grade DNAse I (Sigma-Aldrich). RNA was used to generate cDNA with the First-Strand III cDNA synthesis kit (Invitrogen). Amplification reactions used the PowerUp™ SYBR™ Green Master Mix (ThermoFisher) and the QuantStudio™ 7 Flex Real-Time PCR System (Applied Biosystems).

### Analysis of RNA-seq data

The assembly of *S. pneumoniae* RMV8_rare_ (accession code OX244288) [47] was annotated with Prokka [85]. RNA-seq data (accession codes in Table S4) were mapped to the 2,132 annotated protein coding sequences with Kallisto version 0.46.2 [86] and analysed with Sleuth version 0.30.0 [87]. The patterns of gene transcription in *S. pneumoniae* RMV8_domi_ were used as the reference against which transcription in the other three genotypes (RMV8_rare_, RMV8_domi_ *tvr*::*cat* and RMV8_rare_ *tvr*::*cat*) were compared using Wald tests. The threshold for significant differences in transcription was set at a false discovery rate of 10^-3^ using Q-Q plots (Fig. S25). Data were plotted using the R package circlize [88].

To calculate coverage of each strand of the genome, RNA-seq data were mapped to the whole RMV8_rare_ chromosome with BWA version 0.5.9 [89]. The resulting SAM file was edited by identifying all reverse reads that mapped in a proper pair, and inverting the strand to which they aligned. The coverage of each strand of the genome was then calculated using SAMtools version 1.9 [90].

### Analysis of quantitative PCR data

The ΔΔCt method was used to quantify gene expression through qRT-PCR. The *rpoA* gene was used as a reference gene in each sample, against which the expression of the gene of interest was normalised as a ΔCt value. In all experiments, three technical measurements were recorded for each biological replicate. The mean ΔCt for a selected standard gene in a particular sample was then calculated across technical replicates. This was used to calculate the ΔΔCt values for other genes within the same biological replicate. The fold difference between genotypes was then quantified as 2^-ΔΔCt^. Where possible, RMV8_rare_ was used as the standard sample.

For the quantification of prophage dynamics, the products of primers A and B (“AB”) and A and D (“AD”) were generated from genomic DNA and their concentrations measured using the Qubit Broad Range kit and a Qubit 4 Fluorometer (ThermoFisher). The Ct values for known concentrations of these products were used to generate a standard curve, which was used to convert Ct values from experiments into absolute copy numbers of DNA molecules.

### Assaying transformation efficiency

For assaying induced competence, *S. pneumoniae* genotypes were grown statically in mixed media to an OD_600_ of 0.2-0.25. A 1 mL sample of the bacterial culture was mixed with 5 µL 5 mM CaCl_2_ (Sigma-Aldrich), 100 ng of the *rpoB*_S482F_ PCR product in 2 µL water, and 2.5 µL of the appropriate competence stimulating peptide (Sigma-Aldrich). Tubes were incubated at 35 °C in 5% CO_2_ for 2 h prior to transfer onto solid media. Titrations on non-selective plates were used to calculate the overall number of viable cells. Samples were also spread on agar plates supplemented with 4 µg mL^-1^ rifampicin (Fisher Scientific).

For assaying spontaneous competence, 2×10^5^ cells of RMV8_rare_ were grown in 1 mL mixed media in a 12 well plate at 35 °C, in a 5% CO_2_ atmosphere, with 5 μL 500 mM CaCl_2_ and 1 μg genomic DNA containing a rifampicin resistance marker. Cells were transferred to solid selective media after 16 h of incubation.

### Construction of mutants

The disruption of genes was achieved through amplifying the flanking ∼1 kb regions with primers that added restriction sites on the internal sides of each. An antibiotic resistance cassette was then amplified with the corresponding restriction sites on either side. PCR amplicons were purified following gel electrophoresis as described above, then digested with the appropriate restriction enzyme (*Bam*HI or *Eco*RV; Promega). Both flanking regions were ligated to the resistance marker using T4 DNA ligase (Invitrogen). Transformation was then undertaken as described when using the rifampicin resistance marker.

Transformants were selected on solid media supplemented with the appropriate antibiotic. When using a chloramphenicol acetyltransferase (*cat*) marker, media were supplemented with chloramphenicol (Sigma-Aldrich) at 4 μg mL^-1^. When selecting for isolates carrying the selectable and counter-selectable Janus cassette, media were supplemented with kanamycin (Sigma-Aldrich) at 600 μg mL^-1^; when selecting for isolates lacking the cassette, media were supplemented with streptomycin (Sigma-Aldrich) at 200 μg mL^-1^.

The Janus cassette was used to remove the *cps* locus of R6 Δ*ivr* to generate *S. pneumoniae* R6 Δ*ivr cps*::Janus. This intermediate genotype was transformed with genomic DNA from *S. pneumoniae* 99-4038, followed by selection on streptomycin, to identify bacteria in which the Janus cassette had been lost. Those bacteria which had replaced the cassette with the *cps* locus of 99-4038 were further screened for expression of the capsule, based on their colony morphology.

A modified version of the Janus cassette [91] was constructed in which *rpsL*, causing sensitivity to streptomycin, was replaced with a *pheS*\* sequence, which caused sensitivity to *p*-chloro-phenylalanine (*p*-Cl-Phe) [92]. This Janus*_pheS_* cassette enabled more effective counter-selection against bacteria carrying the cassette using *p*-Cl-Phe (VWR) at 3 μg mL^-1^. For the construction of RMV8_rare_ and RMV8_domi_ *ccnC* and *ccnCL*, the ϕRMV8 prophage and adjacent *ccnCL* gene were replaced with this modified Janus cassette to generate ϕRMV8::Janus*_pheS_* mutants. To restore the original *ccnC* sequence, the relevant locus was directly amplified from *S. pneumoniae* R6. To introduce the modified *ccnCL* without the associated prophage, the *ccnCL* sequence from RMV8 was amplified and ligated to the bacterial sequence flanking the ϕRMV8 *attR* site. Both resulting DNA amplicons were used to transform the ϕRMV8::Janus*_pheS_* mutants, with selection on *p*-Cl-Phe used to identify transformants that had replaced the Janus*_pheS_* cassette. Sequencing was used to confirm the acquisition of the relevant *ccnC* and *ccnCL* alleles.

### Characterisation of the *S. pneumoniae* R6 *cps*_99-4038_ recombinant

Sequencing libraries were generated with the Oxford Nanopore Rapid Barcoding Sequencing Kit (Oxford Nanopore Technologies, code SQK-RBK004), according to the manufacturer’s instructions. Sequencing used a SpotON flow cell installed on a MinION device, generating 304 Mb of data, with a maximum read length of 63,258 bp. Bases were recalled with the high accuracy model of guppy version 6.0.1. Reads were assembled with dragonflye version 1.0.13 [93] using Flye version 2.9 [94] and racon version 2.24 [95], yielding a single 2.03 Mb contig.

The assembly of *S. pneumoniae* R6 *cps*_99-4038_ was aligned to the genome of *S. pneumoniae* R6 using lastz version 1.04.15 [96]. This alignment was filtered to single-fold coverage of the R6 genome with single_cov2 version 11 from the multiz package [97]. The resulting multiple alignment format output was used to generate a FASTA alignment with maf2fasta version 3 [97]. Recombinations were then identified using the pairwise mode of Gubbins v3.2.1 [64].

### Microscopy of pneumococci

Light microscopy of colonies used a Leica DFC3000 G microscope. To prepare samples for transmission electron microscopy, single colonies of pneumococci were frozen to −196 °C at a pressure of 2,100,000 hPa in a few milliseconds using a Bal-Tec HPM 010 machine. Freeze substitution used a Leica EM AFS2 freeze substitution device. The first substitution media was a freshly-made solution of 0.1% tannic acid in acetone with 0.5% glutaraldehyde at −90 °C. After 72 h, samples were washed three times in cold acetone over 2 h. Samples were then infiltrated with 2% osmium tetroxide with 1% uranyl acetate in acetone at −80 °C for 2 h. The temperature was then raised to −20 °C for 16 h, then 4 ℃ for 4 h. The samples were rinsed three times in acetone at room temperature, with further incubation in acetone to dehydrate the samples. Samples were then infiltrated with increasing concentrations of epon in acetone: 30% epon for 4 h; 50% epon overnight; 70% epon for 4 h, 90% for 2 h, then finally two treatments with 100% epon. The samples were then polymerised in fresh epon at 60 °C for 48 h. A diamond knife and a Leica UC6 ultramicrotome were used to cut 50 nm sections, which were mounted on uncoated copper grids. Images were recorded on an FEI 120 keV Biotwin transmission electron microscope with a Tietz F4.16 Charge-Coupled Device.

## Acknowledgements

We thank the Bespoke team at the Wellcome Sanger Institute for generating the sequencing libraries.

## Data availability

Accession codes for the genomic data used to analyse the epidemiology of GPSC12 are listed in Table S1. Accession codes for the genomic data used in the experimental analyses are listed in Table S7. Accession codes for the RNA-seq data are listed in Table S4.

