## Supplementary Figures for "Post-vaccine epidemiology of serotype 3 pneumococci identifies transformation inhibition through prophage-driven alteration of a non-coding RNA"

**Supplementary Materials**

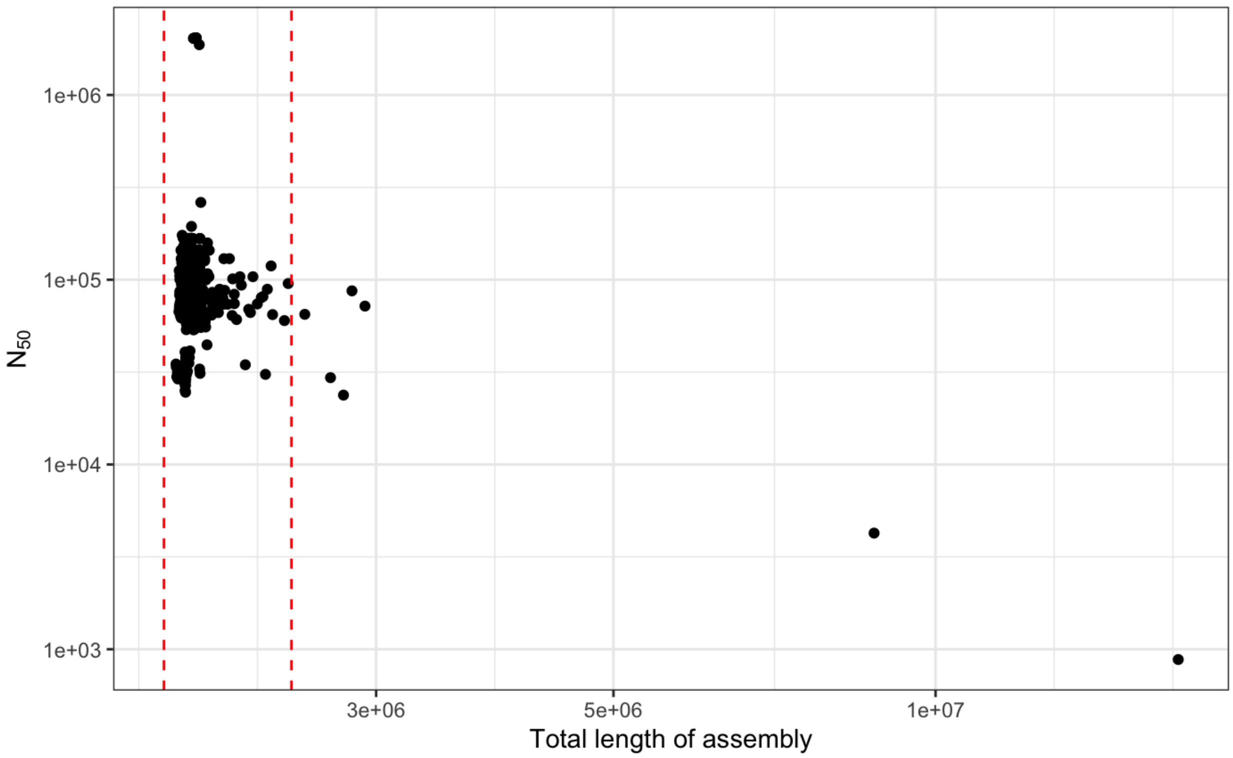


**Figure S1** Exclusion of low-quality datasets. The scatterplot compares the length and $N_{50}$ (the length of the contig spanning the midpoint of a draft assembly, when contigs are arranged in length order) of 978 assemblies generated with SPAdes. The vertical red dashed lines show the threshold distances of 1.9 Mb and 2.5 Mb that were used to identify anomalously long or short sequences. This resulted in the exclusion of seven assemblies from the dataset.

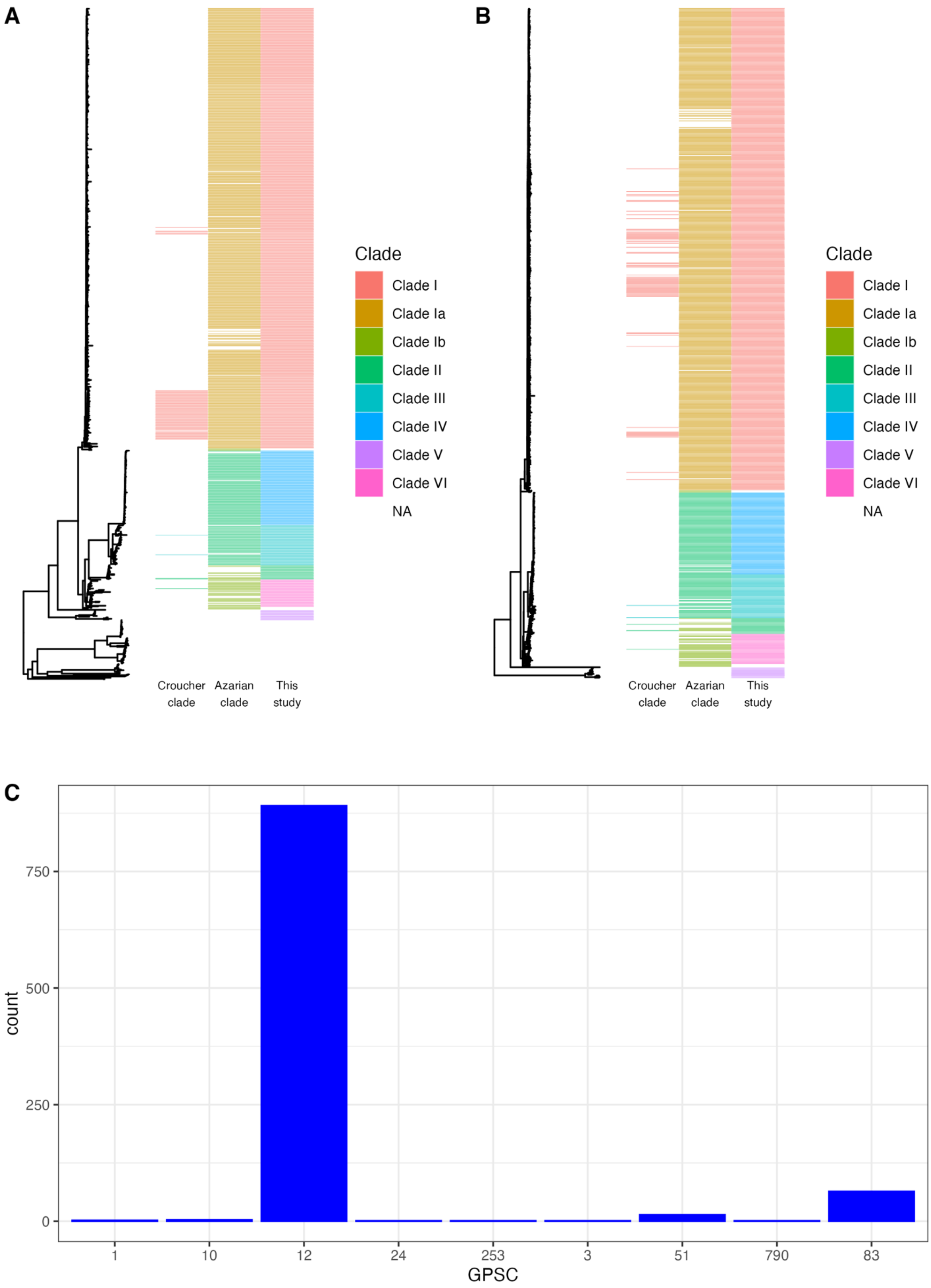
A
B
Clade
Clade I
Clade Ia
Clade Ib
Clade II
Clade III
Clade IV
Clade V
Clade VI
NA
Croucher clade
Azarian clade
This study
C
count
0
250
500
750
1
10
12
24
253
3
51
790
83
GPSC

**Figure S2** Overview of the serotype 3 population. (A) Neighbour-joining tree calculated from core genome distances estimated using PopPUNK across the full serotype 3 dataset of 971 isolates. The columns show the assignment of GPSC12 isolates to clades in Croucher *et al* (2012); Azarian *et al* (2016), which was subsequently extended by Groves *et al* (2019); and this work. (B) Recombination-corrected maximum likelihood phylogeny of 891 GPSC12 isolates. The assignment to clades is shown as in panel A. (C) Bar chart showing the assignment of 971 serotype 3 isolates to GPSCs using PopPUNK and version 6 of the GPS strain database. The majority (891) isolates belonged to GPSC12.

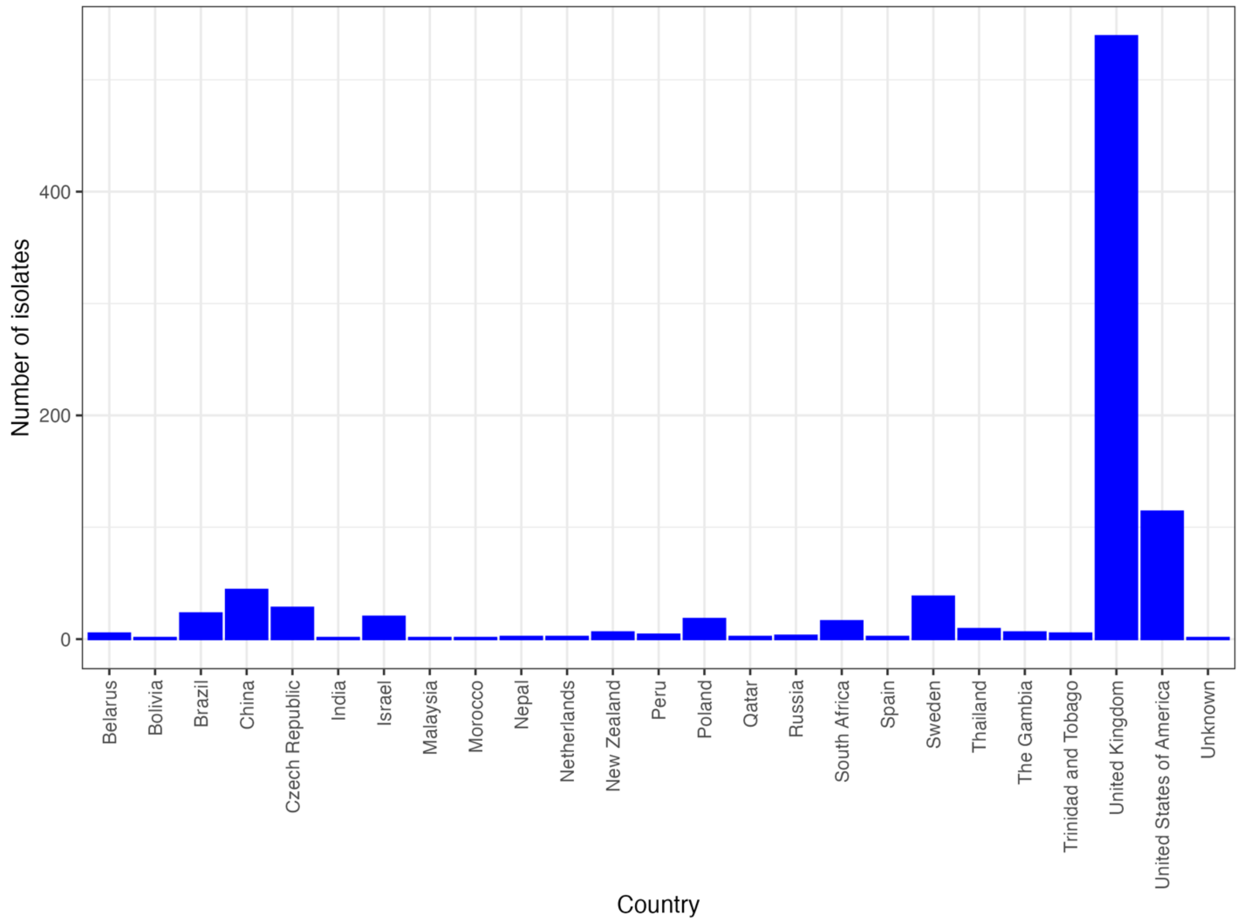


**Figure S3** Geographic distribution of the 891 GPSC12 isolates.

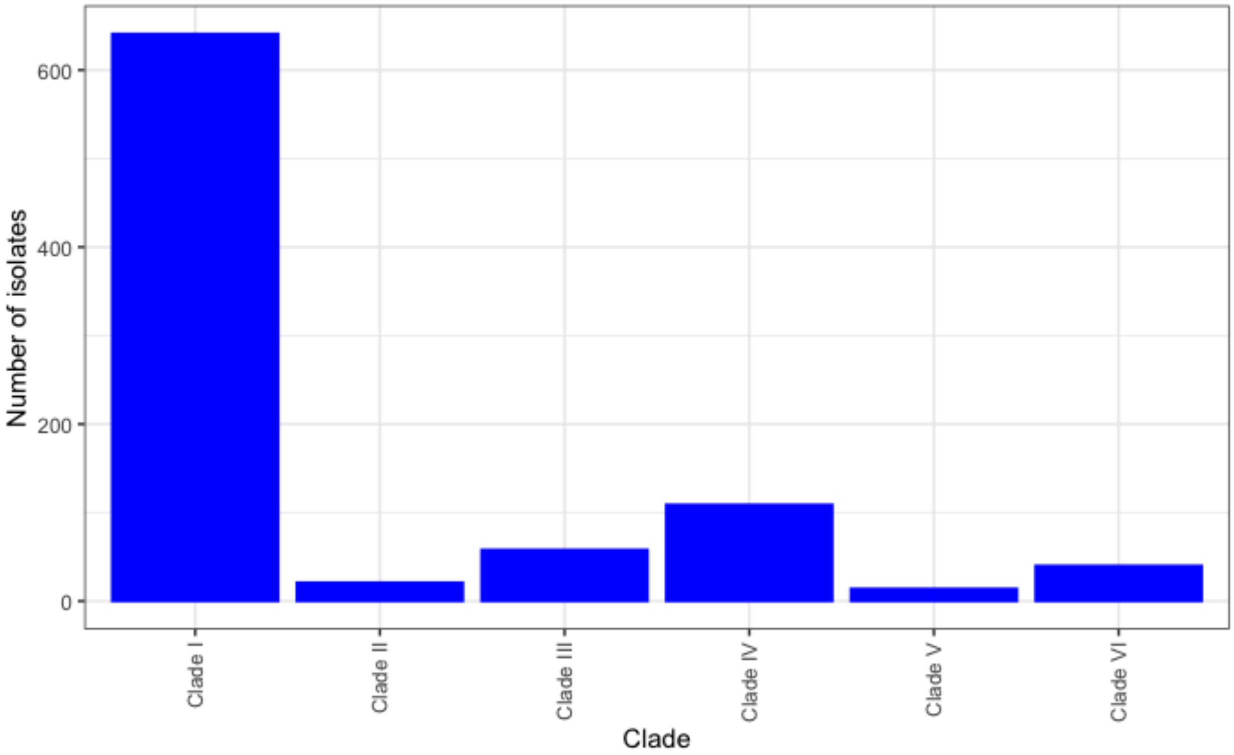


**Figure S5** Distribution of the 891 GPSC12 isolates between the clades defined using the recombination-corrected phylogeny.

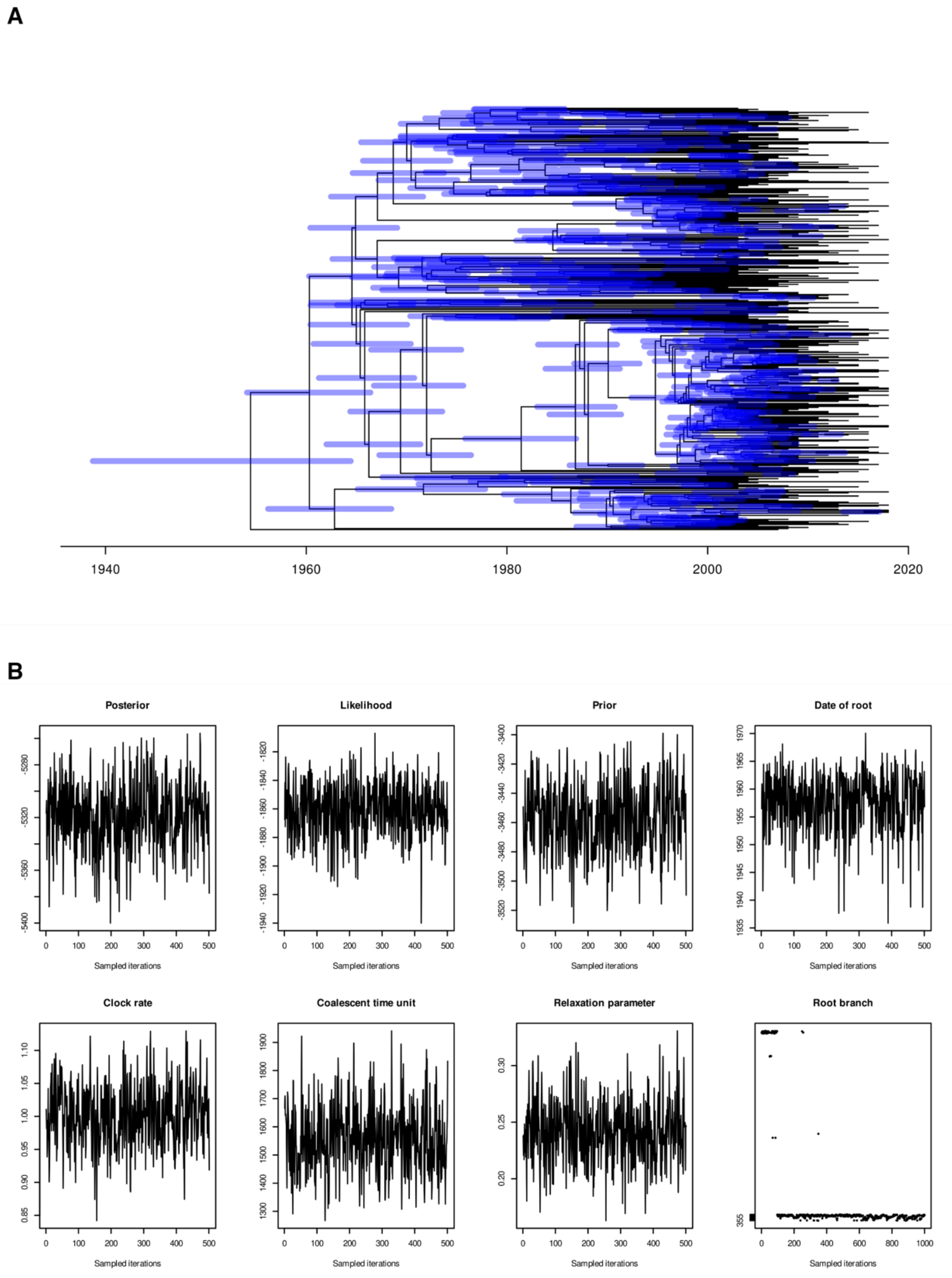


**Figure S4** Phylodynamic analysis of GPSC12 Clade I. (A) Time-calibrated phylogeny of GPSC12 Clade I. The horizontal axis relates the tree to dates. The blue horizontal bars show the 95% credibility intervals for the date of each node in the phylogeny. (B) Markov Chain Monte Carlo sampling of BactDating model parameters over $2.5 \times 10^7$ iterations, after half the chains were removed as warm up. These demonstrate the convergence of the algorithm on a stable set of parameter estimates.

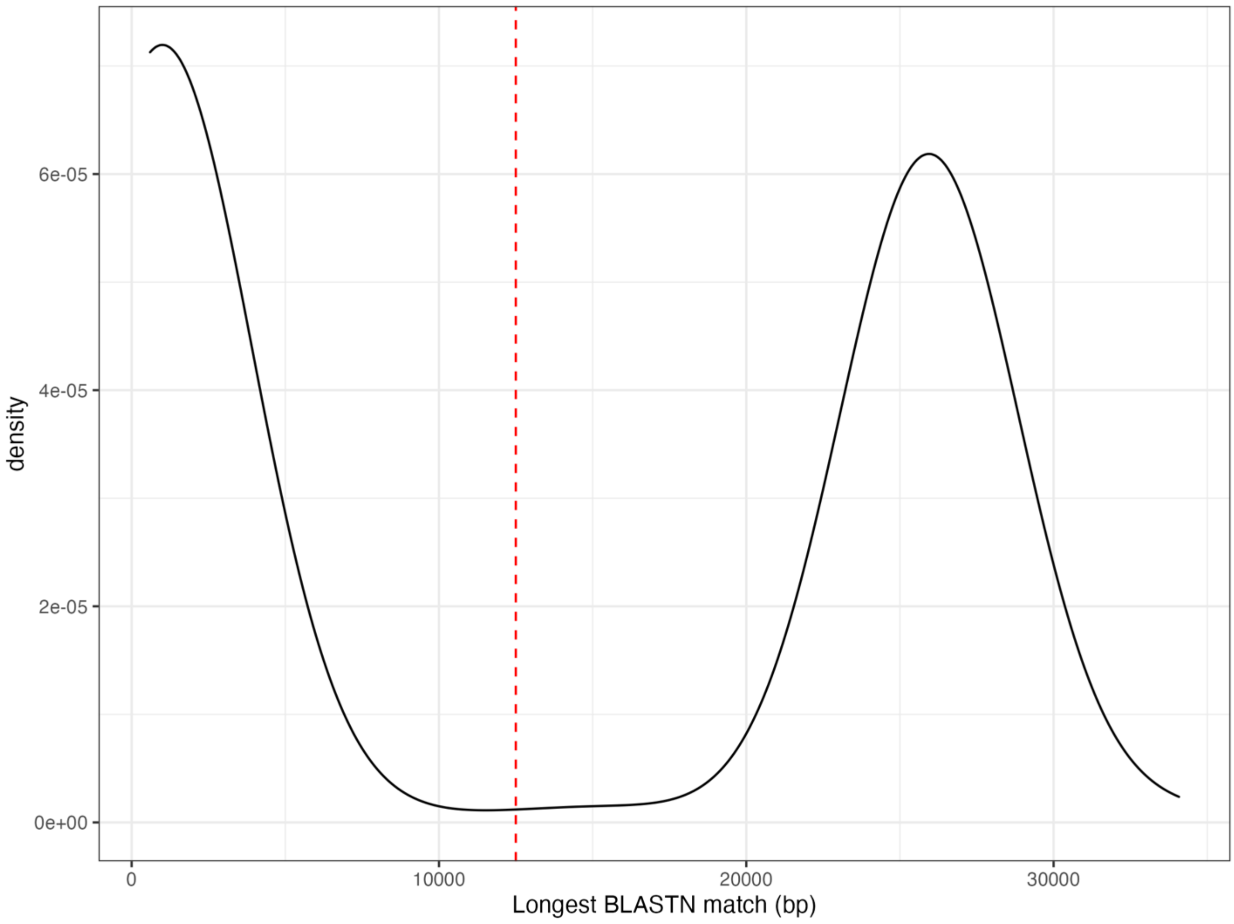


**Figure S6** Inferring the distribution of ϕOXC141-like prophage. The 891 GPSC12 isolates were aligned to the 34,080 bp sequence of ϕOXC141, extracted from *S. pneumoniae* OXC141 (accession code FQ312027). The density plot shows the size distribution of the longest BLASTN hit to the query sequence from each isolate. The red dashed line shows the threshold (12.5 kb) used to distinguish hits to ϕOXC141-like prophage to less specific matches to more divergent prophage.

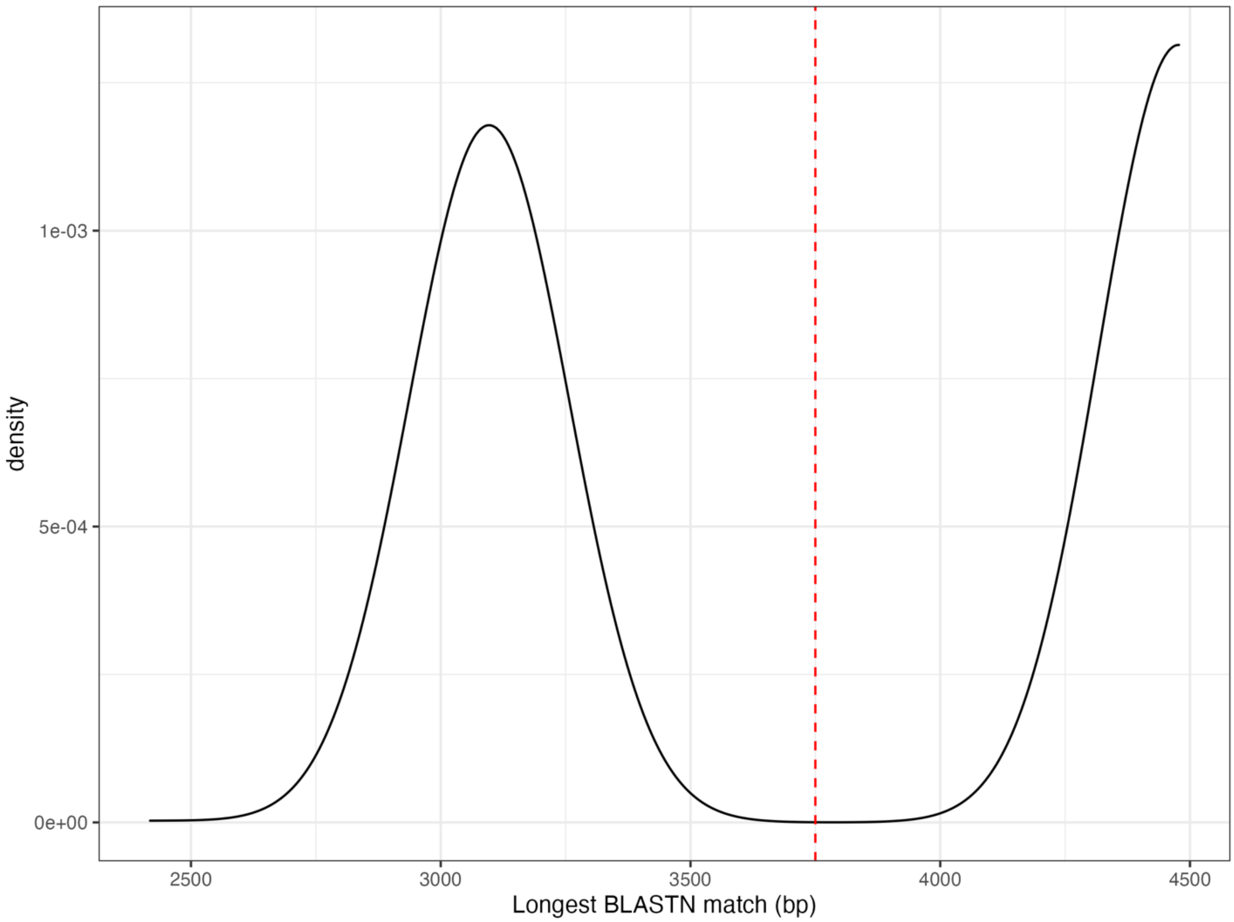


**Figure S7** Inferring the status of the *attB*$_{OXC}$ site. The 891 GPSC12 isolates were aligned to a 4,477 bp sequence spanning the unmodified *attB*$_{OXC}$ insertion site within the *S. pneumoniae* TIGR4 genome (accession code AE005672). The density plot shows the size distribution of the longest BLASTN hit to the query sequence from each isolate. The red dashed line shows the threshold (3,750 bp) used to distinguish the longer, unmodified *attB*$_{OXC}$ sites from those that are shorter, and therefore likely to have been disrupted by a prophage insertion.

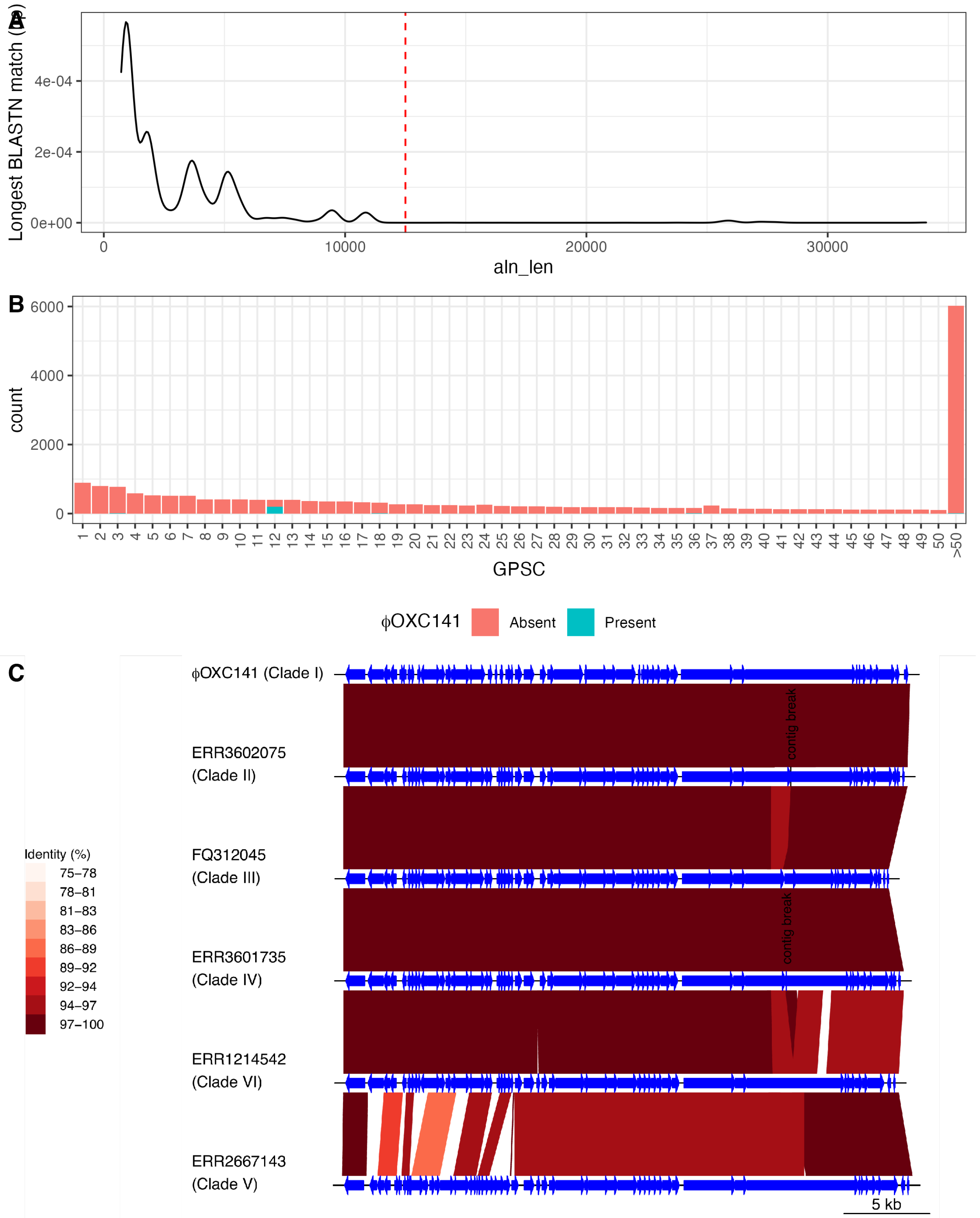


**Figure S8** Identifying the distribution of prophage ϕOXC141. (A) The 34,080 bp sequence of ϕOXC141 was extracted from *S. pneumoniae* OXC141 (accession code FQ312027). The density plot shows the distribution of sizes for the longest BLASTN alignment to each GPS isolate. The red dashed line at 12.5 kb represents the threshold separating hits to full-length ϕOXC141-like prophage from matches to other phage sequences. (B) Distribution of ϕOXC141-like prophage between GPSCs using the 20,047 isolates in the GPS collection. GPSCs with an index above 50 are merged to enable visualisation of the prophage's enrichment in GPSC12. Evidence of ϕOXC141-like prophage was only identified in five isolates outside of GPSC12. (C) Comparison of ϕOXC141-like prophage in GPSC12 isolates outside of Clade I. The blue arrows represent predicted protein coding sequences. The red bands between prophage link regions of similar sequence, with the level of DNA sequence identity indicated by the key. Each prophage is labelled with the accession code of the isolate in which it was identified, and the clade to which the isolate was assigned. Two prophage did not assemble within a single contig, and therefore the contig breaks are marked on the sequences.

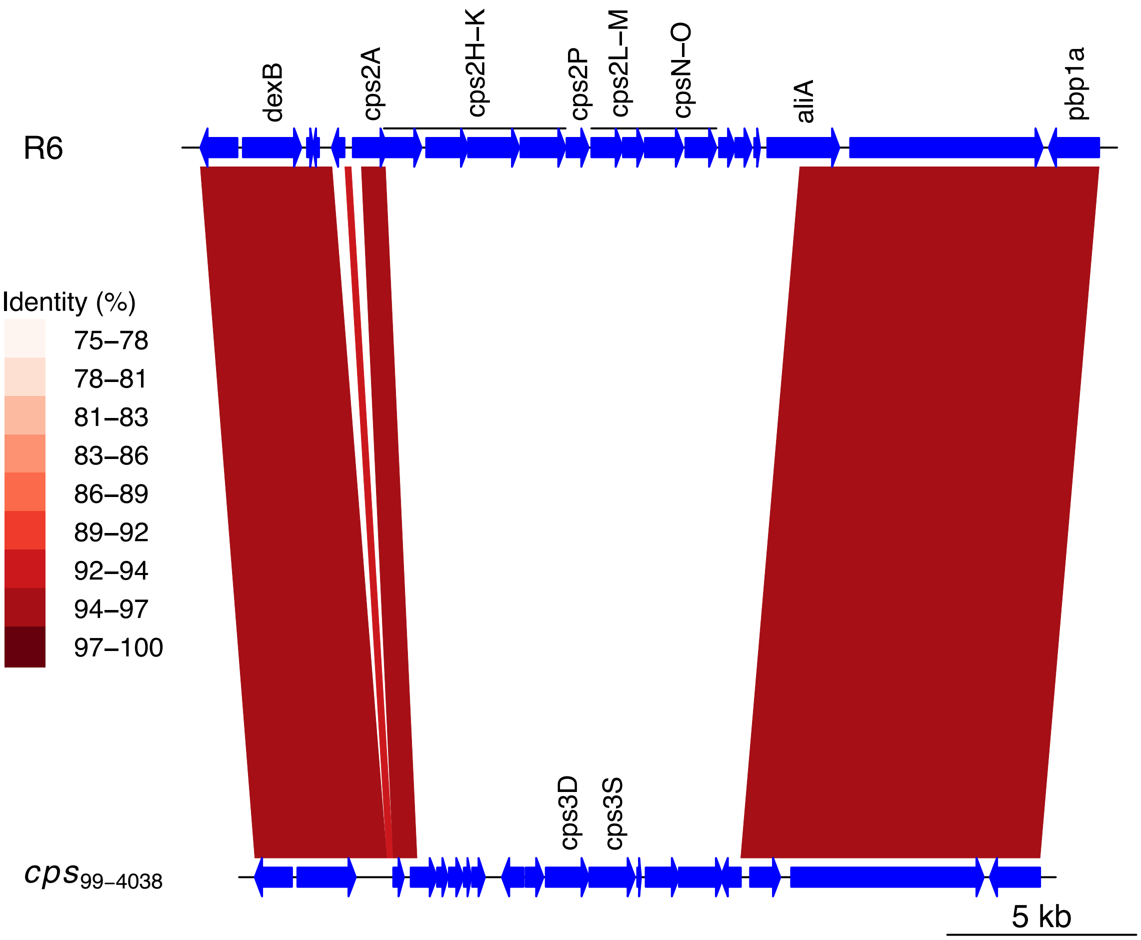


**Figure S9** Replacement of the *cps* locus of *S. pneumoniae* R6 with that from *S. pneumoniae* 99-4038 in the recombinant *S. pneumoniae cps*$_{99\text{-}4038}$. The region encompassed by the recombination inferred to span the *cps* locus in *S. pneumoniae cps*$_{99\text{-}4038}$ is aligned to the orthologous region of the parental genotype, R6, with BLASTN. The red bands link regions of similar sequence, with the colour indicating the level of sequence identity between the pair. The *cps* locus itself is flanked by the *dexB* and *aliA* genes. The serotype 3 allele contains two functional genes: *cps3D*, encoding a UDP-glucose 6-dehydrogenase, and *cps3S*, encoding the polymerase.

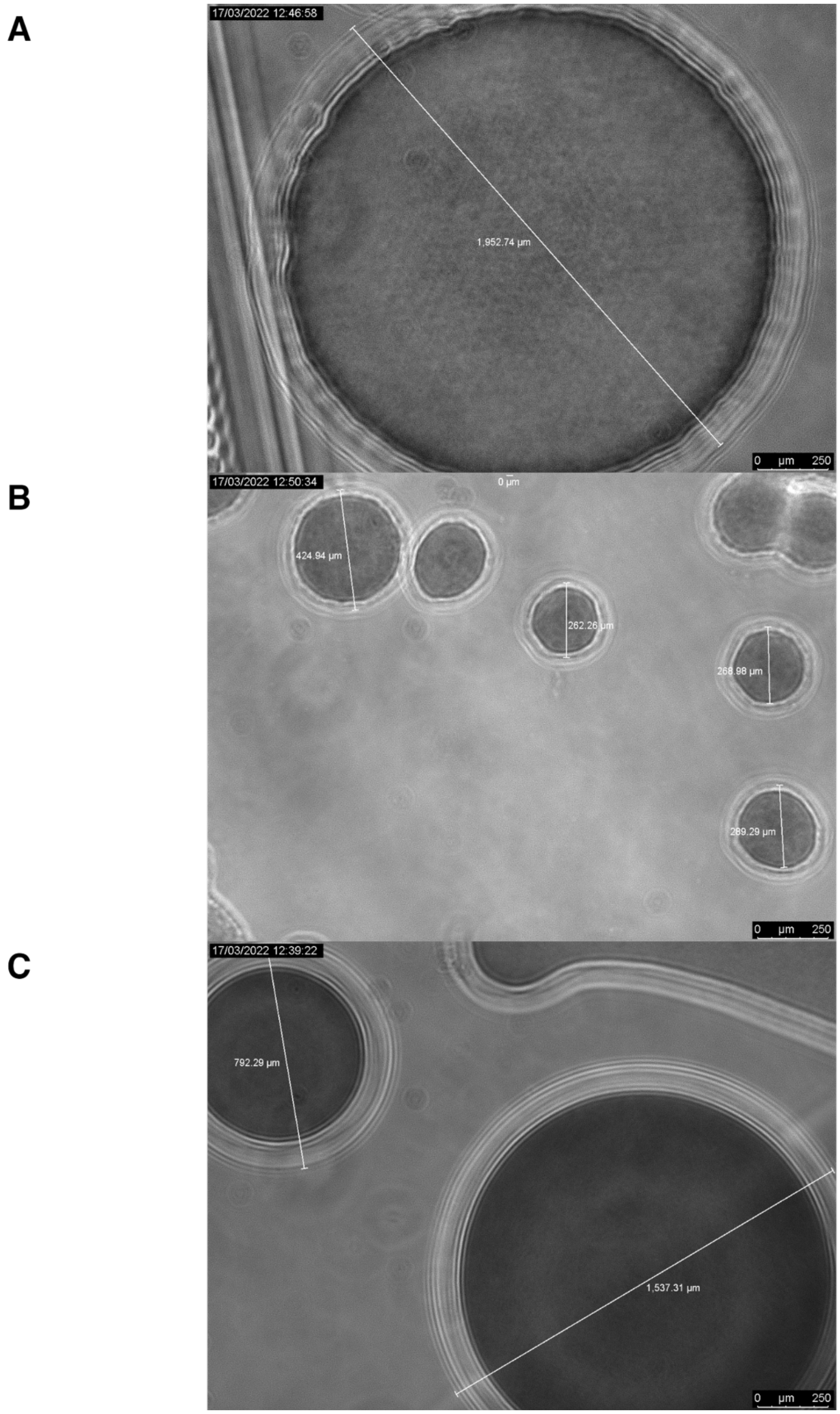


**Figure S10** Light microscopy of pneumococcal colonies. (A) Large mucoid colonies of *S. pneumoniae* 99-4038, a serotype 3 isolate (B) Small colonies of the unencapsulated *S. pneumoniae* R6 (C) Large mucoid colonies of *S. pneumoniae* R6 $cps_{99\text{-}4038}$, engineered to express the serotype 3 capsule.

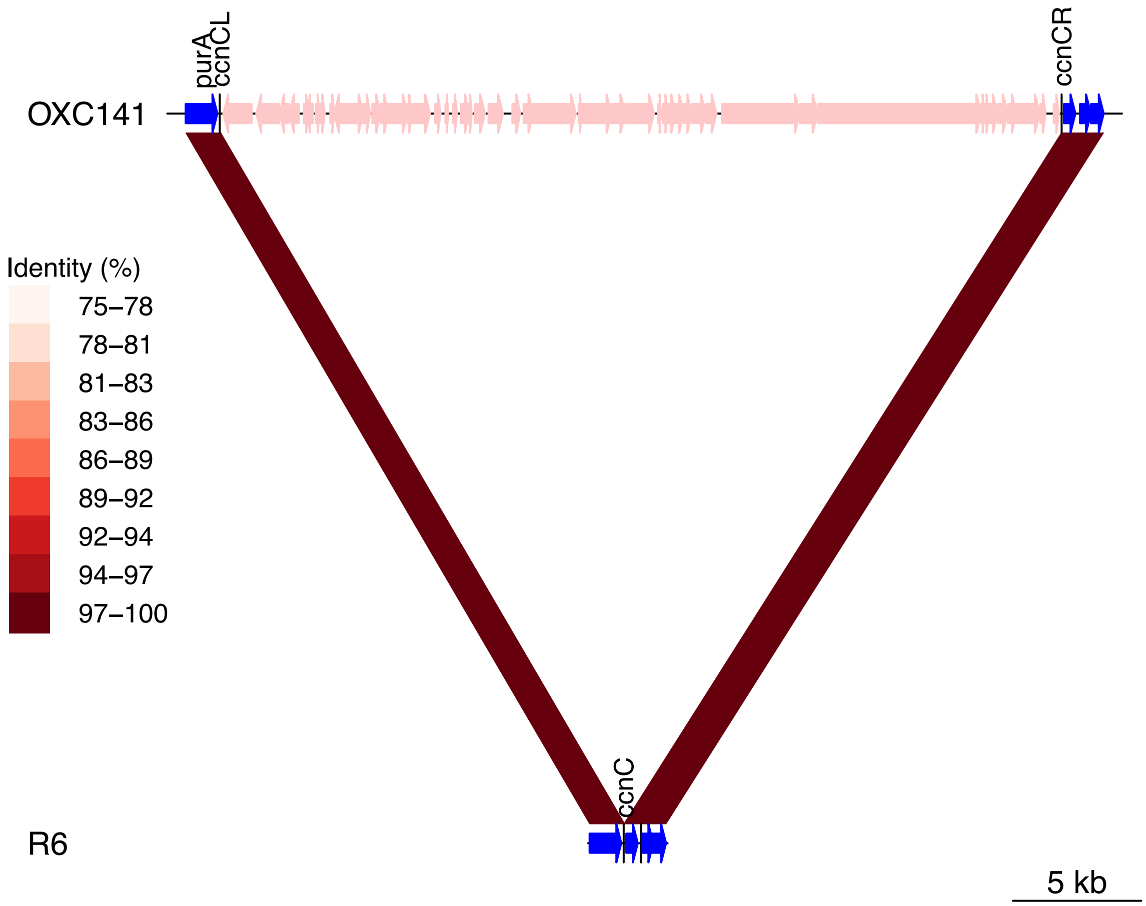


**Figure S11** Modification of *ccnC* through insertion of ϕOXC141 into $attB_{OXC}$. The insertion of ϕOXC141, indicated by the pink genes integrated downstream of *purA*, into the *S. pneumoniae* OXC141 genome is compared to the unmodified $attB_{OXC}$ site of R6 using BLASTN. The red bands link regions of similar sequence, with the colour indicating the level of sequence identity between the pair. The black vertical lines show how the cellular *ccnC* gene at the *attB* site is split into the *ccnCL* and *ccnCR* genes at the *attL* and *attR* sites, respectively.

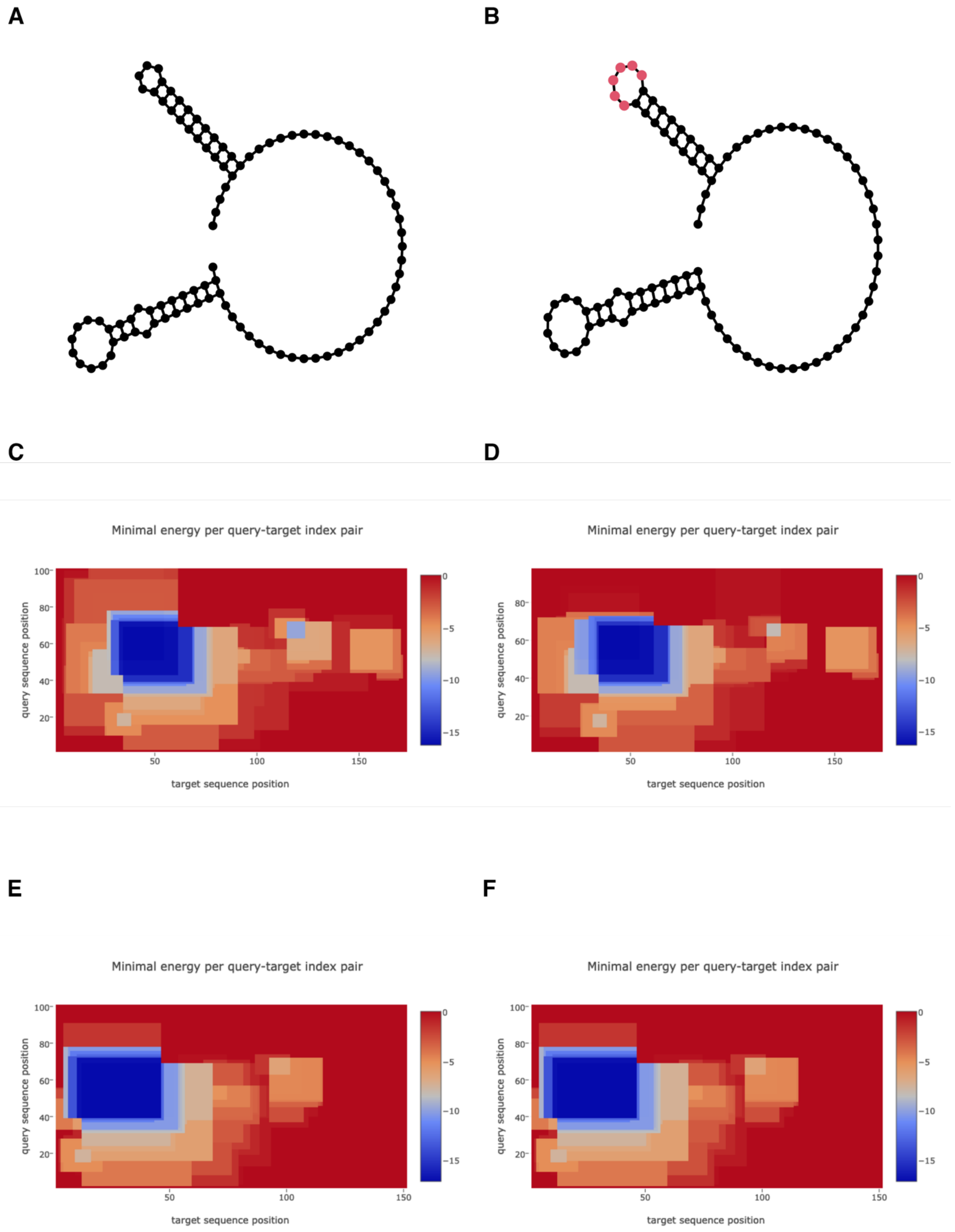
A
B
C
D
Minimal energy per query-target index pair
query sequence position
target sequence position
Minimal energy per query-target index pair
query sequence position
target sequence position
E
F
Minimal energy per query-target index pair
query sequence position
target sequence position
Minimal energy per query-target index pair
query sequence position
target sequence position

**Figure S12** Comparison of the predicted structures of (A) csRNA3 from *S. pneumoniae* R6 and (B) csRNA3L from *S. pneumoniae* OXC141. The sequence "CAAUCA" is highlighted in red. (C) Predicted interaction between csRNA3 from *S. pneumoniae* R6 and the *comC* transcript from *S. pneumoniae* OXC141, which has the CSP1 pherotype. The horizontal axis corresponds to the length of the *comC* mRNA (from the transcription initiation site to the stop codon of the protein coding sequence), and the vertical axis corresponds to the length of csRNA3. The colour shows the minimal energy of the interaction between the sequences at the coordinates specified by the axes in kcal $mol^{-1}$, as indicated by the key. (D) Predicted interaction between csRNA3L from *S. pneumoniae* OXC141 and the *comC* transcript from *S. pneumoniae* OXC141, which has the CSP1 pherotype. Data are displayed as described for panel (C). (E) Predicted interaction between csRNA3 from *S. pneumoniae* R6 and the *comC* transcript from *S. pneumoniae* RMV8, which has the CSP2 pherotype. Data are displayed as described for panel (C). (F) Predicted interaction between csRNA3L from *S. pneumoniae* OXC141 and the *comC* transcript from *S. pneumoniae* RMV8, which has the CSP2 pherotype. Data are displayed as described for panel (C).

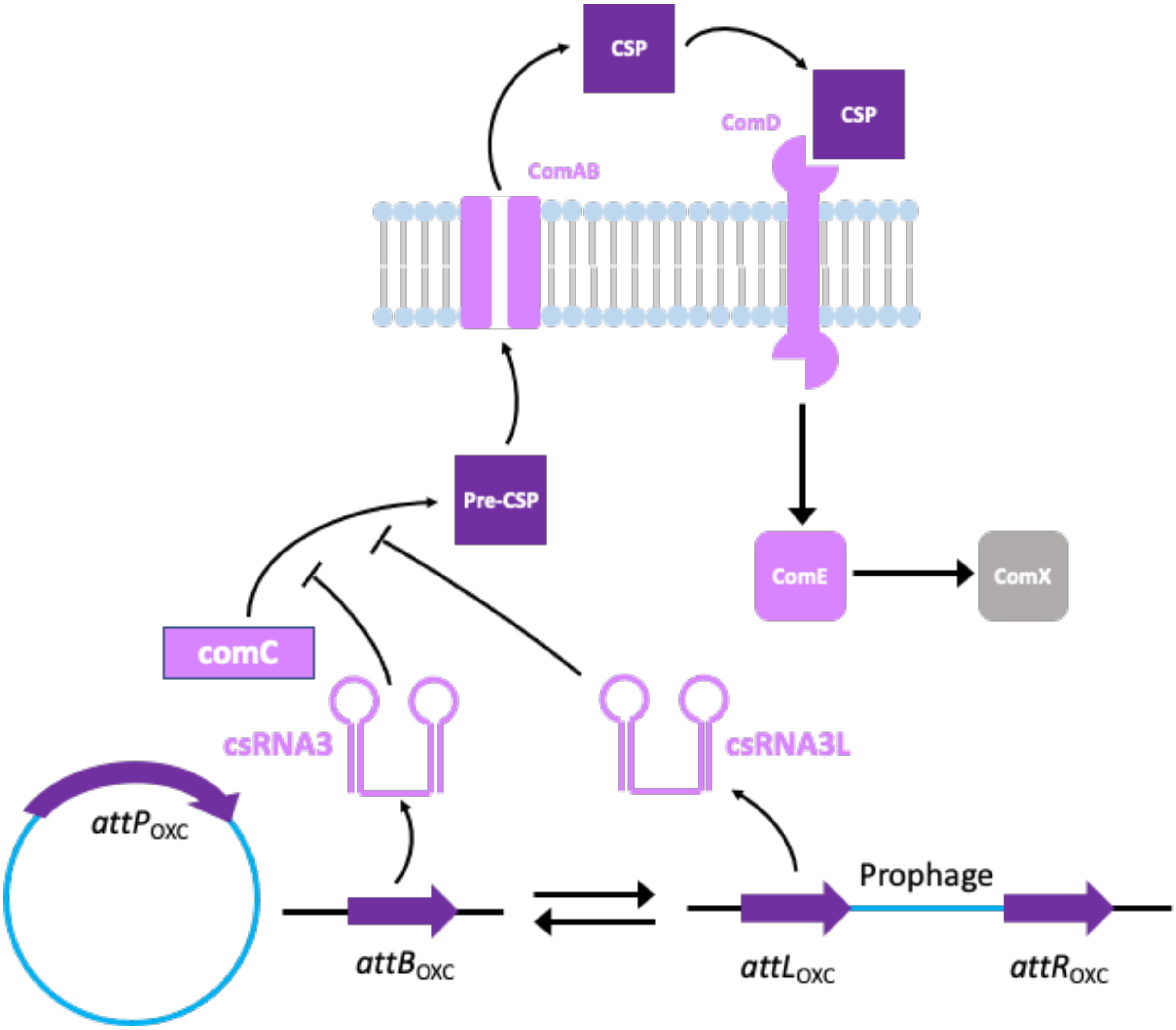


**Figure S13** Model of csRNA3L inhibition of competence. Pneumococcal competence is regulated by a quorum-sensing mechanism. The *comC* gene encodes a pre-peptide that is exported and processed into Competence Stimulating Peptide (CSP) by the ComAB transporter. CSP is detected by the ComDE two competent system, which activates the transcription of early competence genes such as *comX*. The ComX alternative sigma factor then drives expression of the late competence genes required for uptake of DNA from the environment. The csRNA3 RNA reduces the concentration of *comC* transcripts, thereby inhibiting the production of CSP. This delays or prevents the induction of competence by endogenously-produced CSP, but does not affect the induction of competence by exogenously-supplied CSP. The enhanced activity of csRNA3L should therefore further delay or inhibit the spontaneous induction of the competence machinery, without affecting cell responses to CSP added *in vitro*.

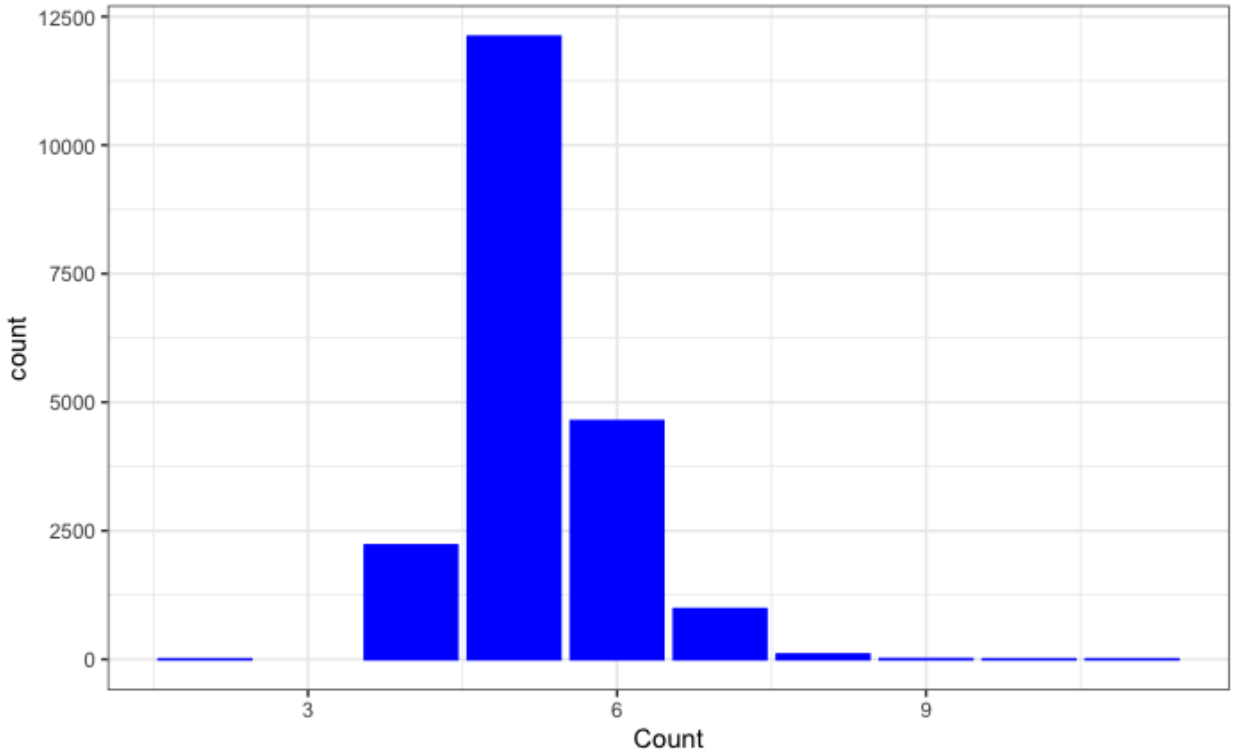


**Figure S14** Distribution of csRNA sequences across GPS isolates. The bar chart shows the number of csRNA sequences identified in each of the 20,047 GPS isolates.

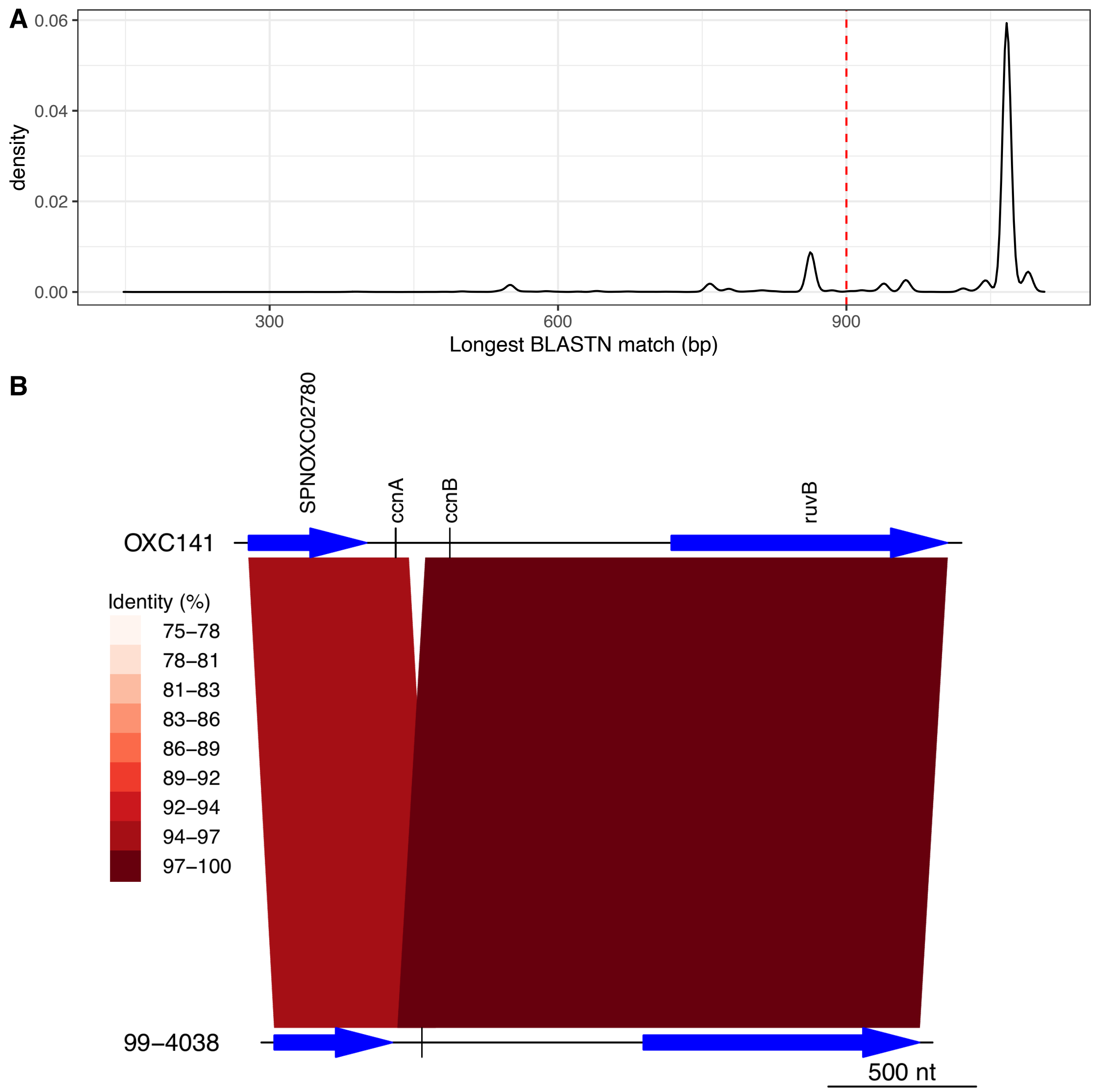


**Figure S15** Identifying deletions removing *ccnA*. (A) A 1,067 bp sequence spanning the region upstream of *ruvB*, which encodes *ccnA* and *ccnB*, was extracted from *S. pneumoniae* R6 and aligned to the 20,047 GPS isolates. The density plot shows the distribution of sizes for the longest BLASTN hit to each GPS isolate. The red dashed line at 900 bp represents the threshold separating the full-length loci from those having undergone a deletion in this part of the genome. (B) Formation of a chimeric *ccnAB* gene through an intragenomic recombination between *ccnA* and *ccnB*. The complete genomes of Clade I isolates *S. pneumoniae* OXC141 and 99-4038 were aligned with BLASTN. The blue arrows show protein coding sequences, and the black vertical lines mark the non-coding csRNA genes. The colours of the bands indicate the level of similarity between the sequences. The V-shape of the bands shows how the tandem csRNA genes at this locus have recombined and deleted the intervening sequence, generating *ccnAB* within *S. pneumoniae* 99-4038.

**A**

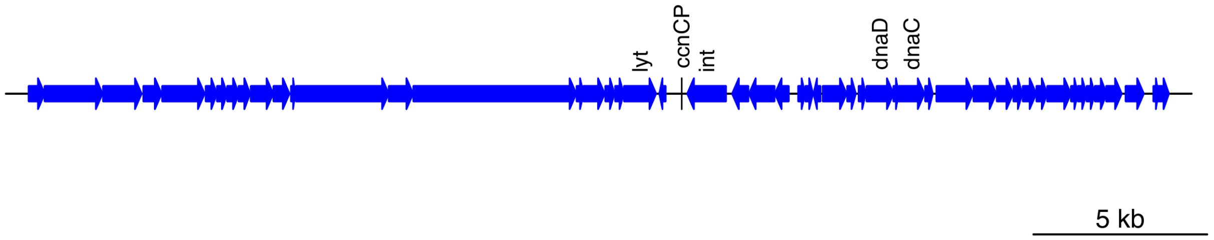


**B**

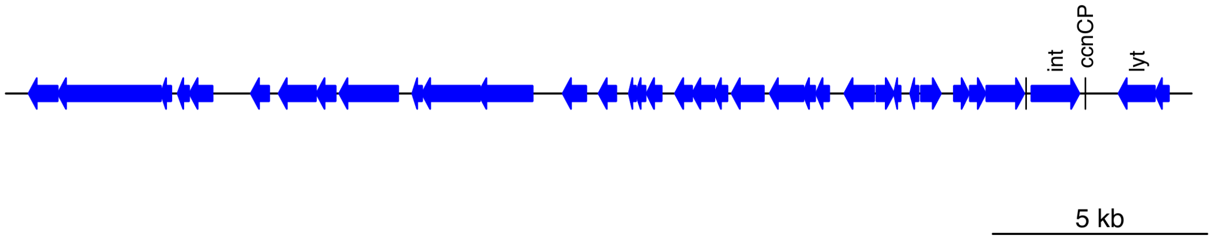


**Figure S16** Identification of *ccnCP* on pneumococcal prophage sequences. (A) Annotation of the prophage *S. pneumoniae* SpSL1 (accession code KM882824), showing the presence of *ccnCP* between the *int* and *lyt* genes, corresponding to the *attP* site of the phage. (B) Annotation of the prophage *S. pneumoniae* ϕARI0578 (accession code KT337360), showing the presence of *ccnCP* between the *int* and *lyt* genes, corresponding to the *attP* site of the phage.

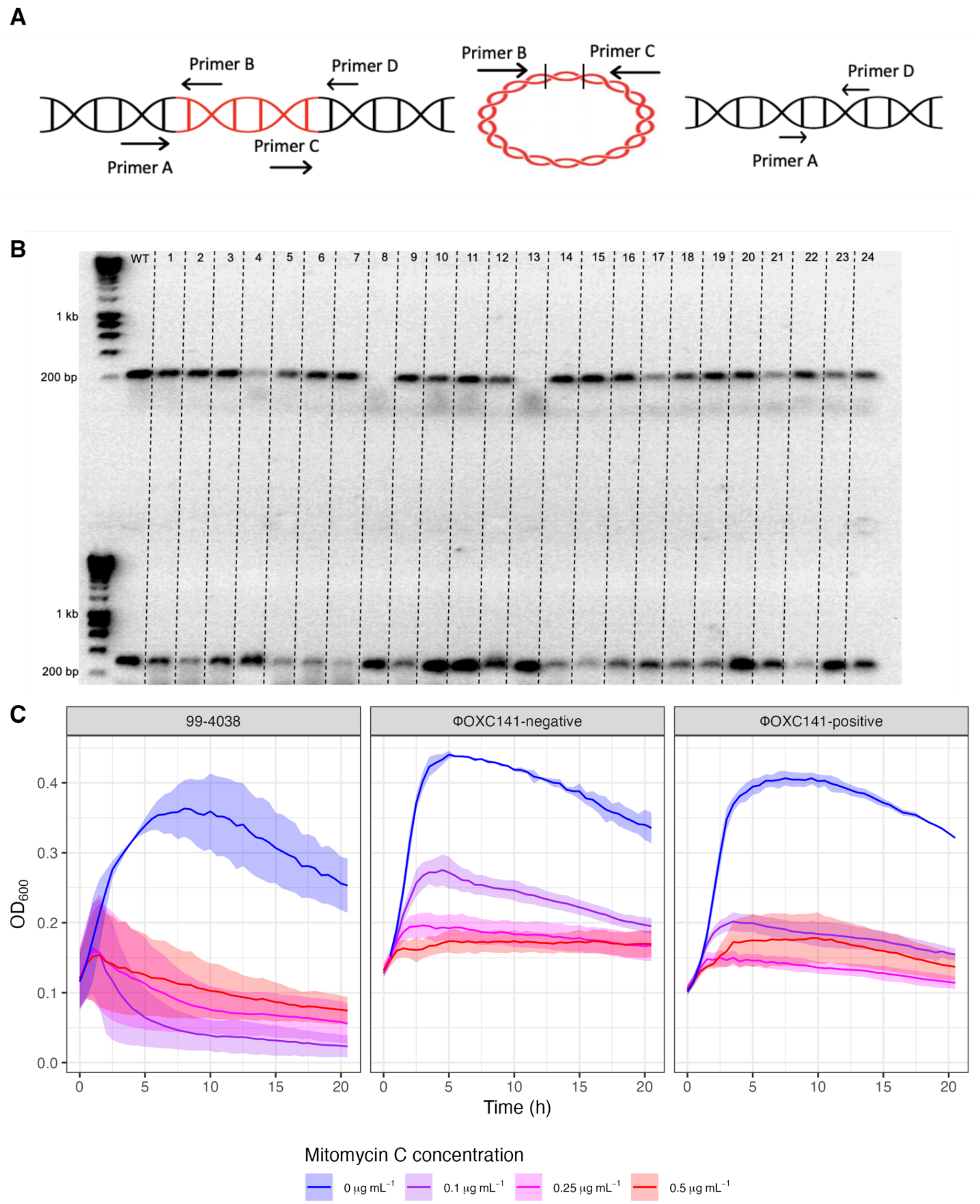


**Figure S17** Isolation of *S. pneumoniae* 99-4038 ΔϕOXC141. (A) Schematic showing how four primers were used to determine the distribution and activity of prophage. Primers A and B were used to detect the $attL_{OXC}$ site of an integrated prophage; primers B and C were used to detect the $attP_{OXC}$ site on circular excised prophage; and primers A and D were used to detect the *attB*$_{OXC}$ site in bacteria in which the prophage has excised or been deleted. (B) Agarose gel showing the A and B, and A and D, PCR products for the wild-type 99-4038 genotype (WT) and 24 colonies isolated following three rounds of mitomycin C exposure in a passage experiment. The size markers are a HyperLadder 1kb (Bioline). The top row shows the products of primers A and B (expected size of 217 bp), indicating the presence of an integrated ϕOXC141. The bottom row shows the products of primers A and D (expected product size of 307 bp), indicating the presence of an intact *attB*$_{OXC}$ site resulting from prophage excision or deletion. Each column corresponds to a different isolate; those numbered 8 and 13 were confirmed to have the ΔϕOXC141 genotype by further PCR experiments. (C) Mitomycin C sensitivity of the parental *S. pneumoniae* 99-4038 isolate, and ϕOXC141-negative and ϕOXC141-positive isolates from the third round of the passage. The isolates found to be ϕOXC141-negative by PCR amplification also exhibited higher tolerance of mitomycin C than those that were found to be ϕOXC141-positive. This would be expected if the prophage has been deleted, rather than excised.

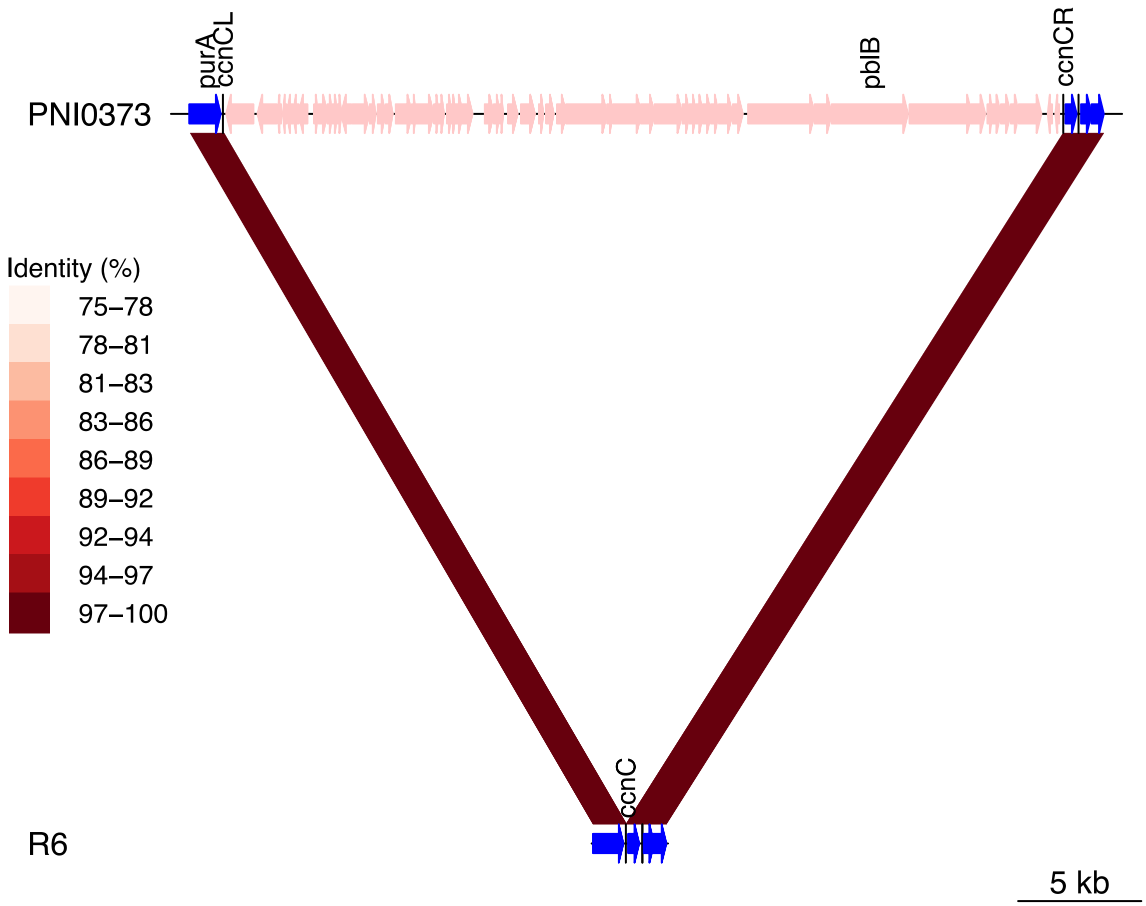


**Figure S18** Modification of *ccnC* through insertion of ϕPNI0373 into *attB*$_{OXC}$. The insertion of ϕPNI0373, indicated by the pink genes integrated downstream of *purA*, into the *S. pneumoniae* PNI0373 genome (accession code CP001845) is compared to the unmodified *attB*$_{OXC}$ site of R6 using BLASTN. The red bands link regions of similar sequence, with the colour indicating the level of sequence identity between the pair. The black vertical lines show how the cellular *ccnC* gene at the *attB* site is split into the *ccnCL* and *ccnCR* genes at the *attL* and *attR* sites, respectively.

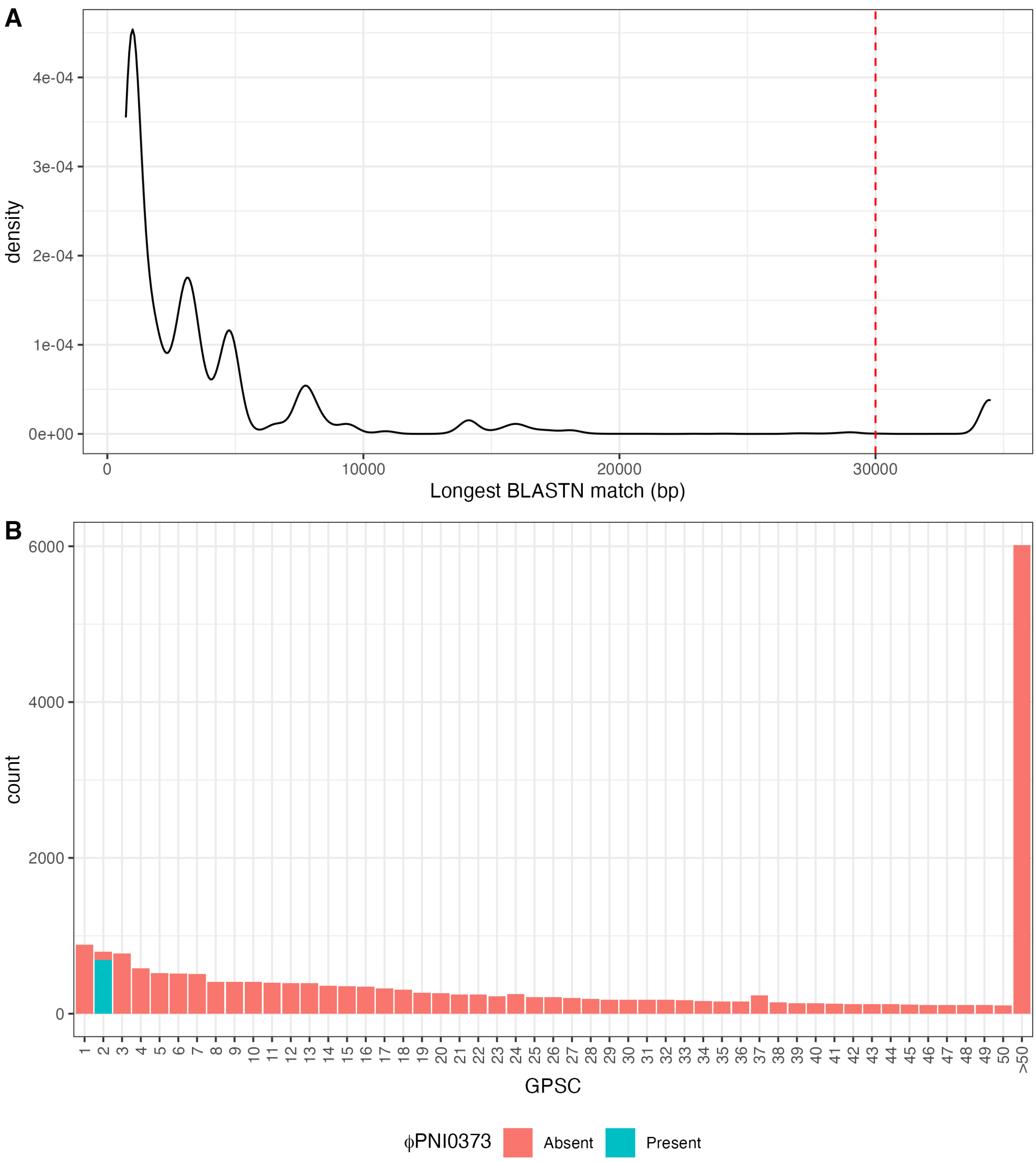


**Figure S19** Identifying the distribution of prophage ϕPNI0373. (A) The 34,437 bp sequence of ϕPNI0373 was extracted from *S. pneumoniae* PNI0373 (accession code CP001845). The density plot shows the distribution of sizes for the longest BLASTN hit to each GPS isolate. The red dashed line at 30 kb represents the threshold separating alignments to full-length ϕPNI0373-like prophage from matches to other phage sequences. (B) Distribution of ϕPNI0373-like prophage between GPSCs using the 20,047 isolates in the GPS collection. GPSCs with an index above 50 are merged to enable visualisation of the prophage's enrichment in GPSC2. No evidence was found of ϕPNI0373-like prophage outside of GPSC2.

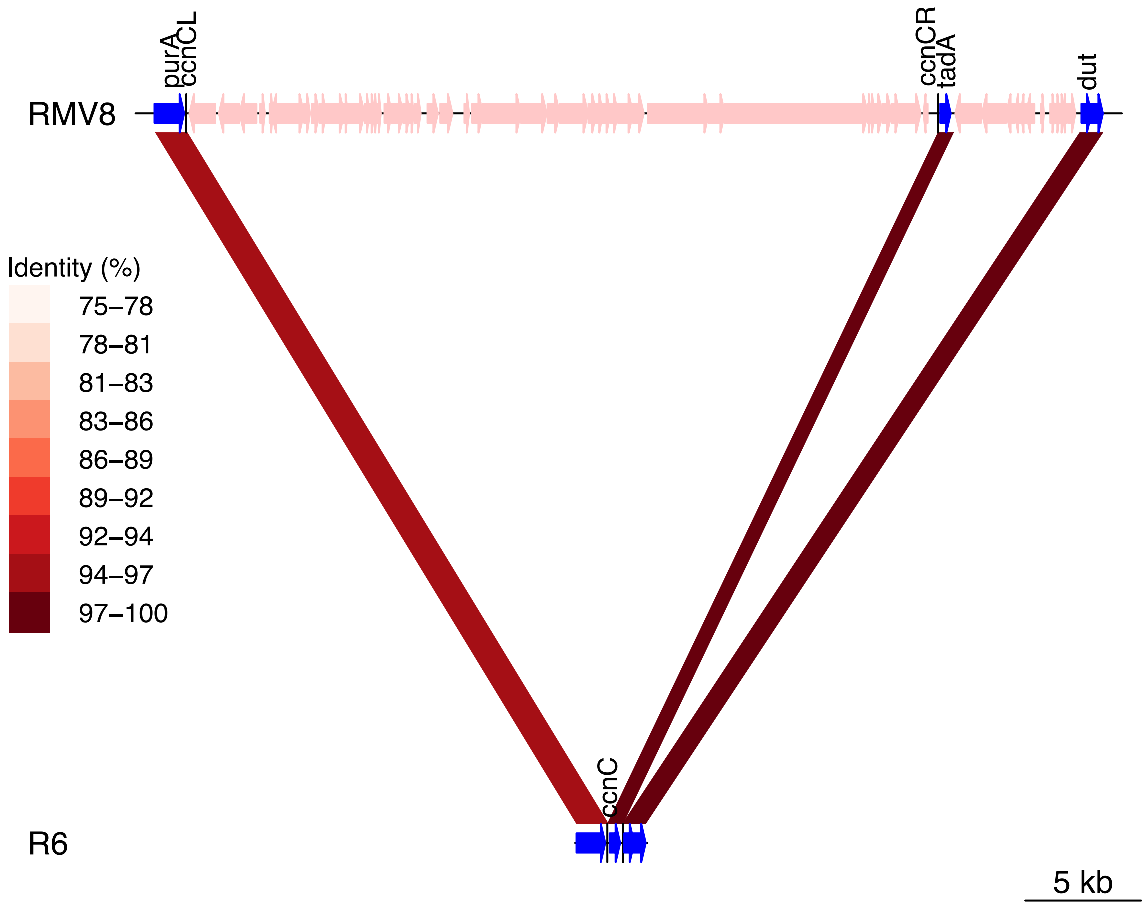


**Figure S20** Modification of *ccnC* through insertion of ϕRMV8 into *attB*$_{OXC}$. The insertion of ϕRMV8, indicated by the pink genes integrated downstream of *purA*, into the *S. pneumoniae* RMV8 genome (accession code OX244288) is compared to the unmodified *attB*$_{OXC}$ site of R6 using BLASTN. The red bands link regions of similar sequence, with the colour indicating the level of sequence identity between the pair. The black vertical lines show how the cellular *ccnC* gene at the *attB* site is split into the *ccnCL* and *ccnCR* genes at the *attL* and *attR* sites, respectively. The second set of pink genes, inserted on the right of ϕRMV8, represents a phage-related chromosomal island (PRCI) inserted adjacent to the *tadA* gene.

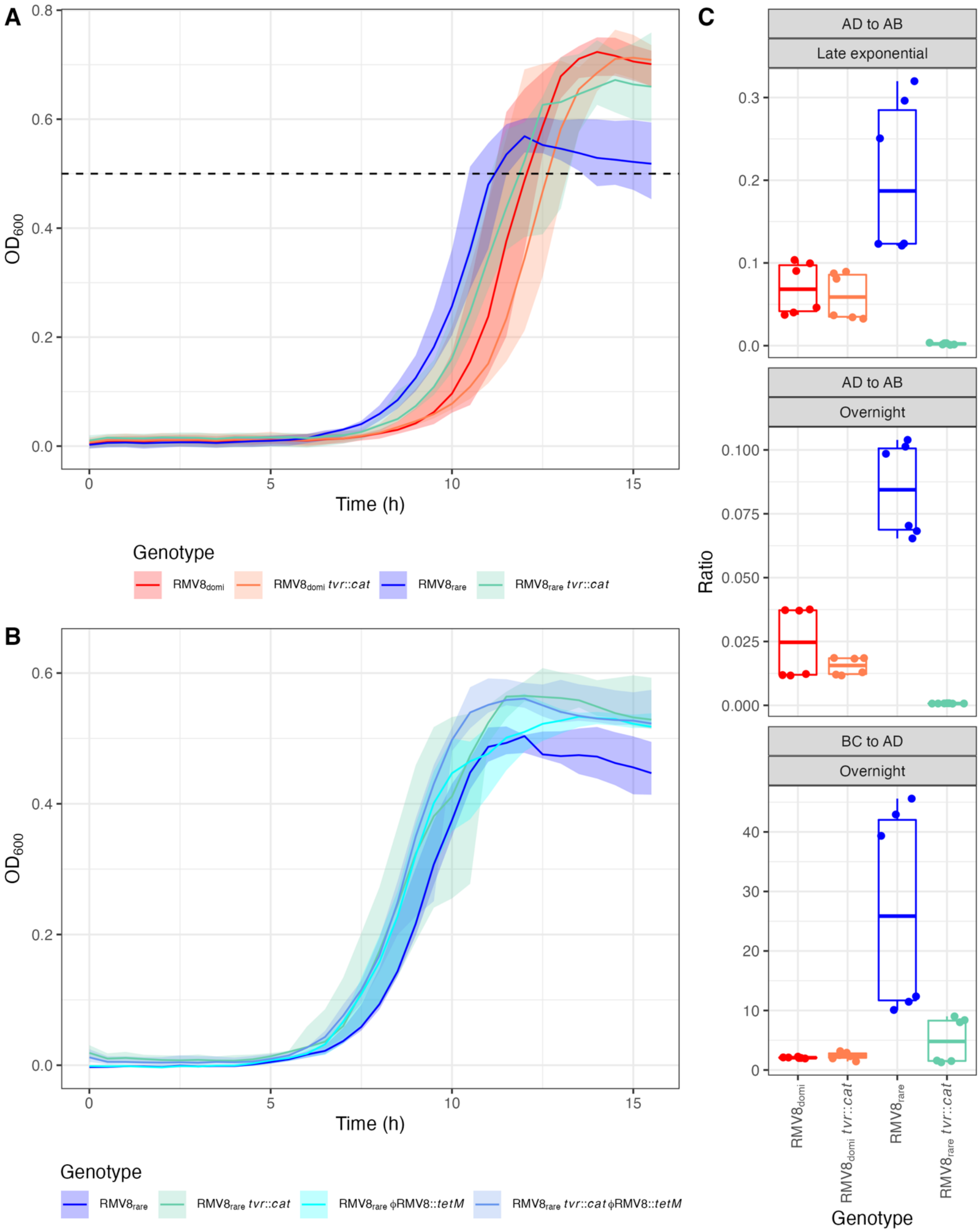


**Figure S21** Growth characteristics of RMV8 genotypes. (A) Growth curves of *S. pneumoniae* RMV8 genotypes analysed by RNA-seq. The solid line shows the median, and the shaded region shows the minimum to maximum range, from eight replicate experiments for each genotype. The atypical growth profile is that of *S. pneumoniae* RMV8$_{rare}$, which has a growth defect in late exponential phase, relative to other genotypes. The dashed horizontal line at an $OD_{600}$ of 0.5 corresponds to the cell density at which samples for RNA-seq were taken for each genotype. (B) Growth curves for $RMV8_{rare}$ genotypes. Data are shown as in panel (A), but summarise four replicate experiments per genotype. The growth defect of $RMV8_{rare}$ is eliminated by the replacement of prophage ϕRMV8 with a *tetM* resistance marker. (C) Assays of the activity of ϕRMV8 using quantitative PCR (qPCR). The same primer arrangements are used as displayed in Fig. S17. The proportion of phage excised from the chromosome was estimated by qPCR of an amplicon spanning the *attL* site (using primers A and B), the *attB* site (using primers A and D), and the *attP* site (using primers B and C). The ratios of the product of primers A and D to the product of primers A and B were used to quantify the level of prophage excision for the four genotypes analysed by RNA-seq during the late exponential phase ($OD_{600}$), and after overnight growth. The ratios of the product of primers B and C to the product of primers A and D were used to quantify the level of circularised phage for the same four RMV8 genotypes after overnight growth.

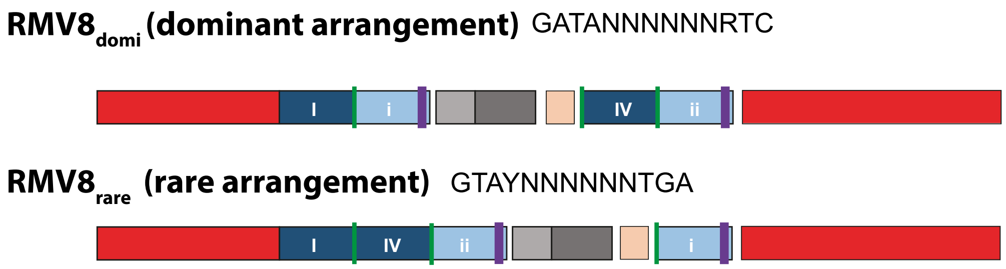


**Figure S22** Differential arrangement of the *tvr* loci distinguishing $RMV8_{domi}$ and $RMV8_{rare}$. Each is labelled with the DNA motif they target. The red boxes correspond to the conserved methylase (*hsdM*, at the 5' end) and endonuclease (*hsdR*, at the 3' end) genes. The blue boxes correspond to the sequences encoding the target-recognition domains (TRDs) of the specificity subunit. The dark blue boxes correspond to the 5' TRD-encoding sequences, and the light blue boxes correspond to the 3' TRD-encoding sequences. The active specificity subunit gene (*hsdS*) is formed by the combination of a 5' and 3' TRD-encoding sequence that is closest to the 5' end of the locus. The HsdS protein determines the motif targeted by the system encoded by the *tvr* locus. The rearrangement of the TRD-encoding sequences is driven by TvrR, a recombinase encoded by a gene represented by the pink box, which is regulated by the products of the *tvrAT* genes, represented by the grey boxes. The *tvrR* gene is truncated in these "locked" variants to limit the interconversion between arrangements at the *tvr* locus.

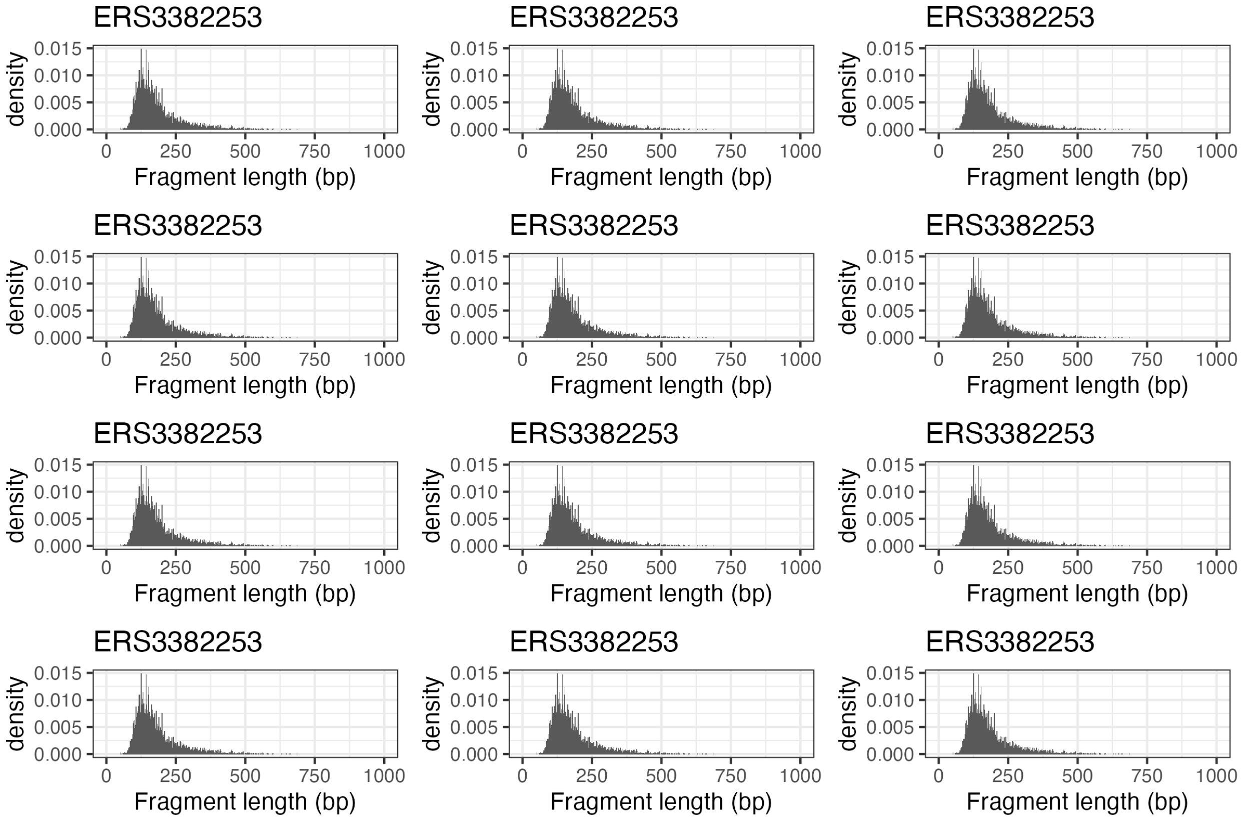


**Figure S23** Bar plots showing the inferred distribution of fragment sizes from mapping by Kallisto. The consistency of these distributions between samples, each labelled with its accession code (Table S4), suggests there should not be any bias introduced by differences in sequencing library preparation.

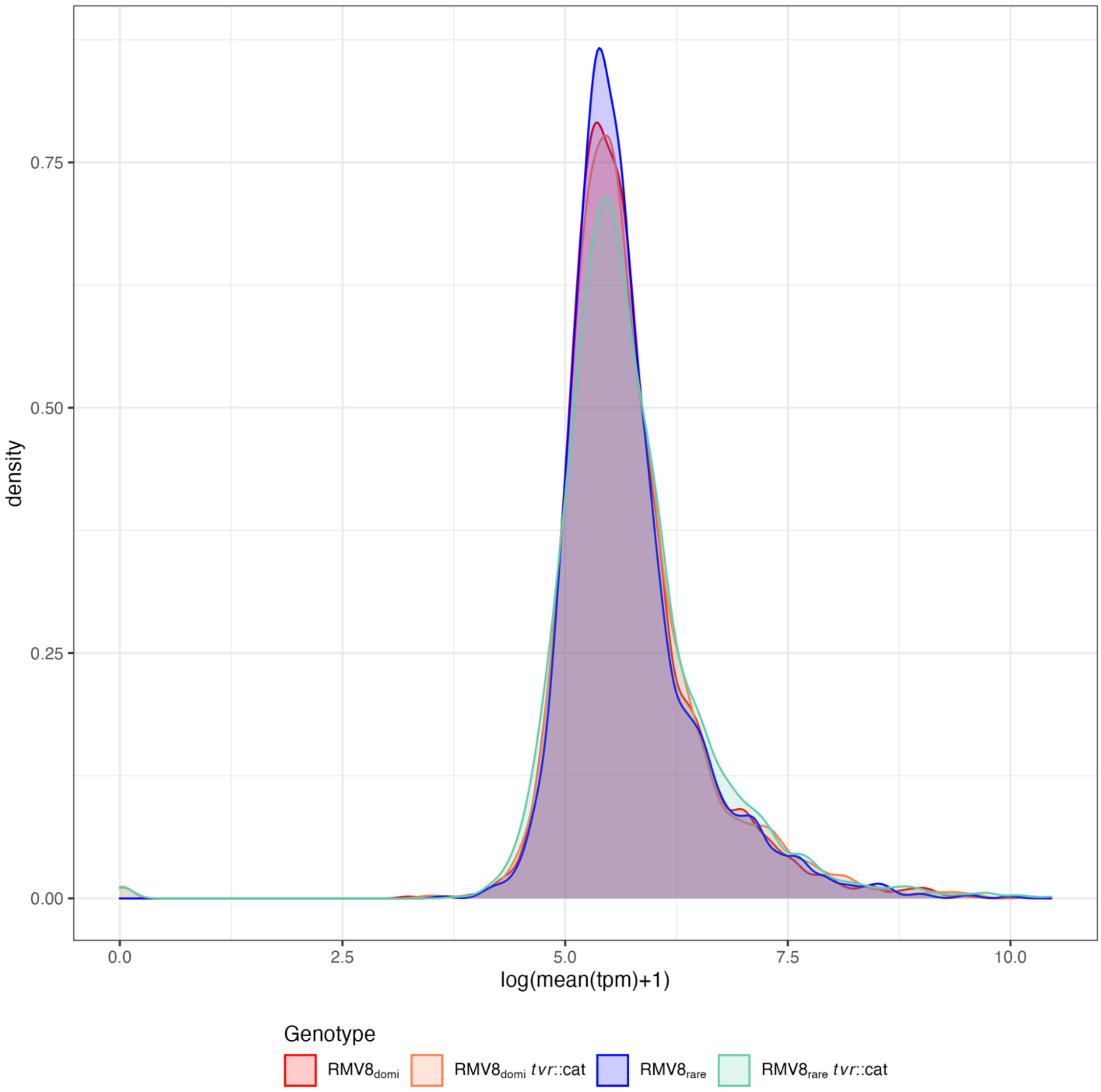


**Figure S24** Density plots showing the distribution of mean transcripts per million (tpm) values across coding sequences for samples in different groups. The consistency of these distributions suggests there should not be any systematic biases causing false positive inferences of transcriptional differences between samples.

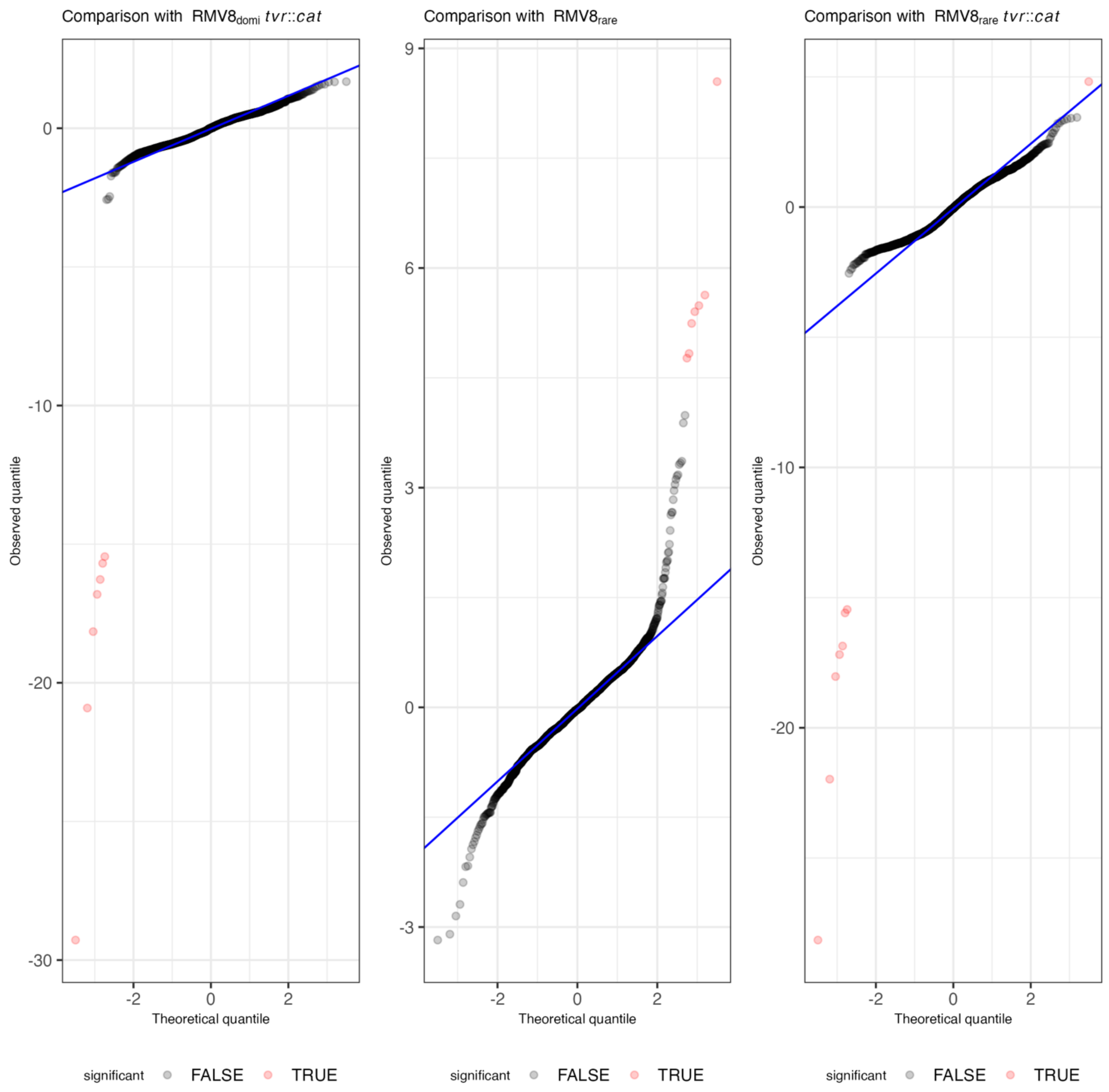


**Figure S25** Q-Q plots comparing the theoretical and observed distributions of the Wald test statistic across genes for the contrast of transcriptional patterns between $RMV8_{domi}$ and the other analysed genotypes, as indicated by the plot titles. The blue line shows the relationship expected under the null hypothesis of no difference in expression patterns. Each point represents a coding sequence. Points are coloured red if the null hypothesis can be rejected at a false discovery rate of $10^{-3}$, following a Benjamini-Hochberg correction for multiple testing. The Q-Q plot shows this threshold captures the major differences between the genotypes.

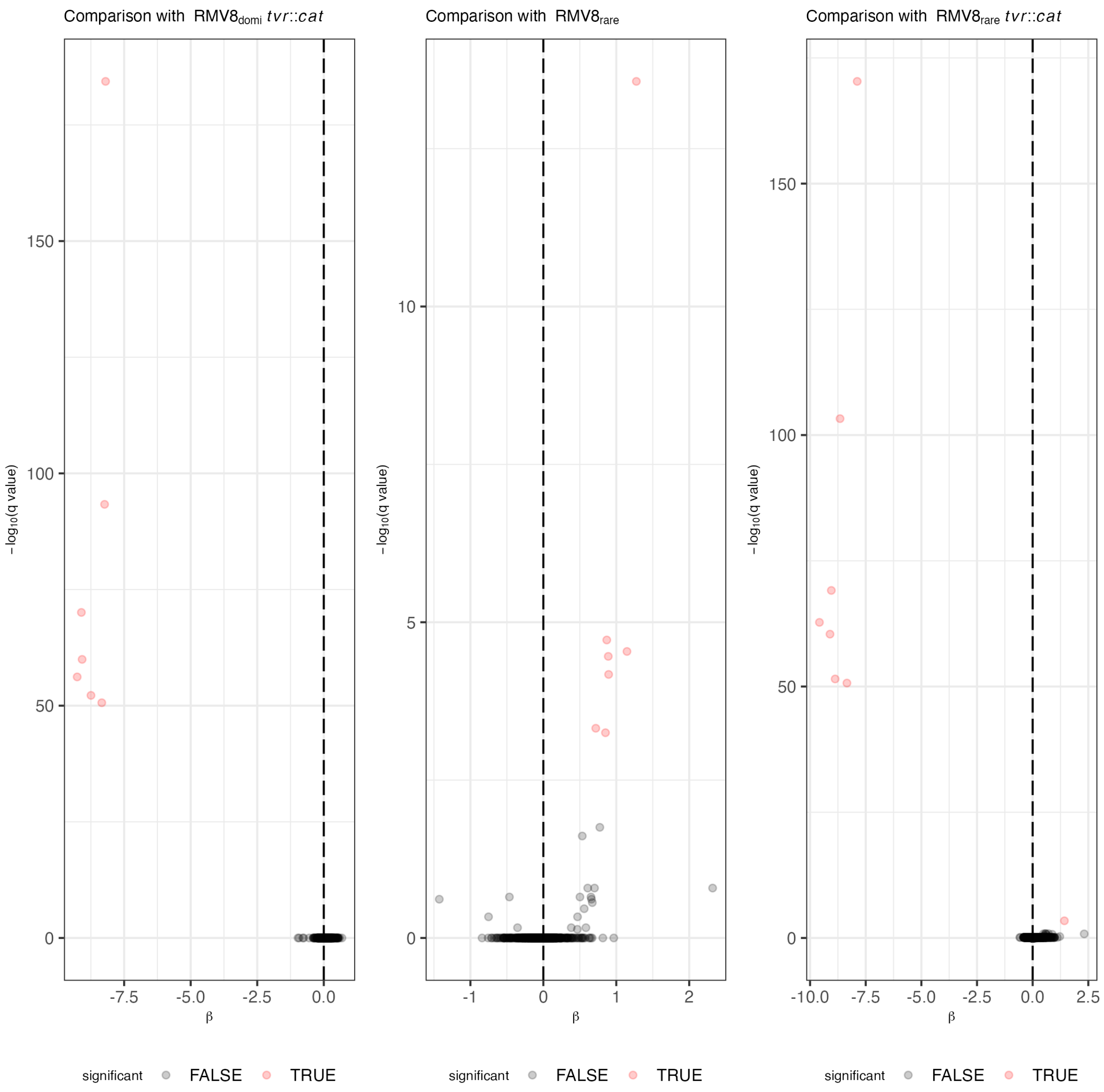


**Figure S26** Volcano plots contrasting the transcriptional patterns between RMV8$_{domi}$ and the other analysed genotypes, as indicated by the plot titles. The horizontal axis shows the natural logarithm of the fold difference in expression levels between the genotypes, β. The vertical axis shows the negative base 10 logarithm of the *q* value following a Benjamini-Hochberg correction. Points are coloured red where this value exceeds the false discovery rate threshold of $10^{-3}$.

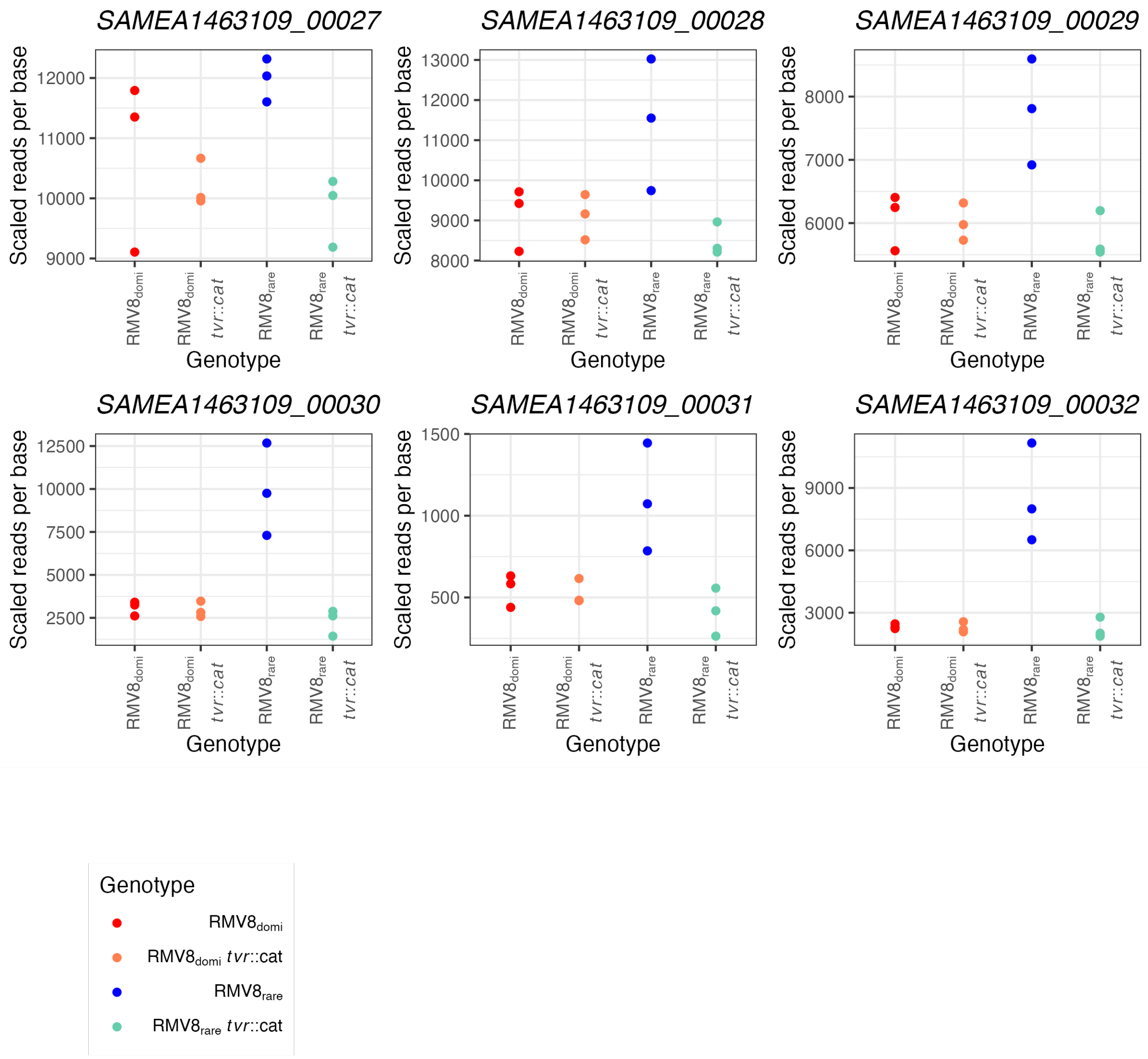


**Figure S27** Quantification of the expression of ϕRMV8 lysogeny genes using RNA-seq data. Points are coloured using the scheme described in Fig. 4, as shown in the key. Each plot shows the transcription of a different gene, in scaled reads per base, across the four genotypes. These genes tend to be most highly expressed in $RMV8_{rare}$.

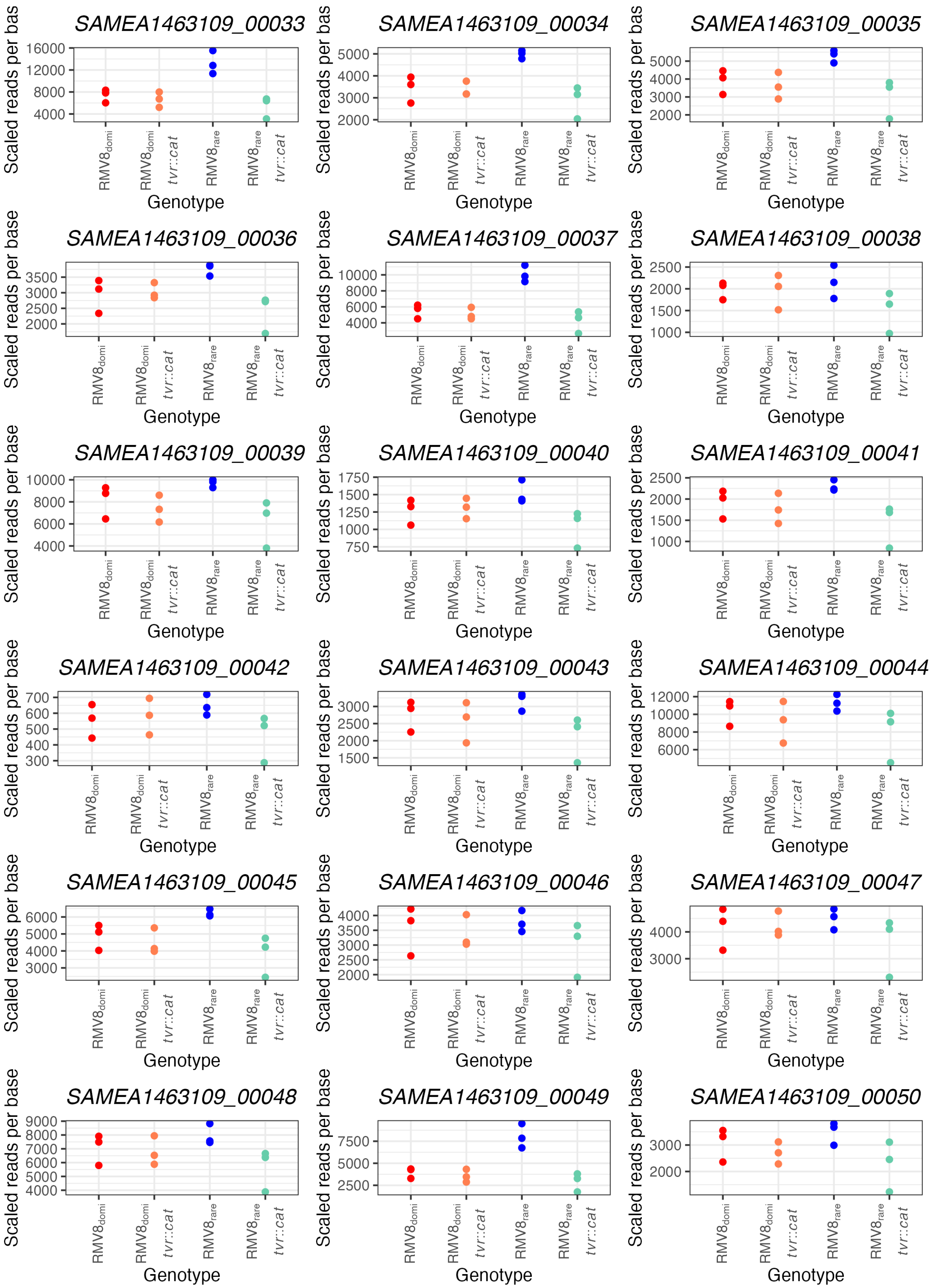
SAMEA1463109_00033
SAMEA1463109_00034
SAMEA1463109_00035
SAMEA1463109_00036
SAMEA1463109_00037
SAMEA1463109_00038
SAMEA1463109_00039
SAMEA1463109_00040
SAMEA1463109_00041
SAMEA1463109_00042
SAMEA1463109_00043
SAMEA1463109_00044
SAMEA1463109_00045
SAMEA1463109_00046
SAMEA1463109_00047
SAMEA1463109_00048
SAMEA1463109_00049
SAMEA1463109_00050
Scaled reads per base
Genotype
RMV8domi
RMV8domi tvr::cat
RMV8rare
RMV8rare tvr::cat

**Figure S28** Quantification of the expression of ϕRMV8 replication genes using RNA-seq data. Points are coloured using the scheme described in Fig. 4. Each plot shows the transcription of a different gene, in scaled reads per base, across the four genotypes. These genes tend to be most highly expressed in $RMV8_{rare}$.

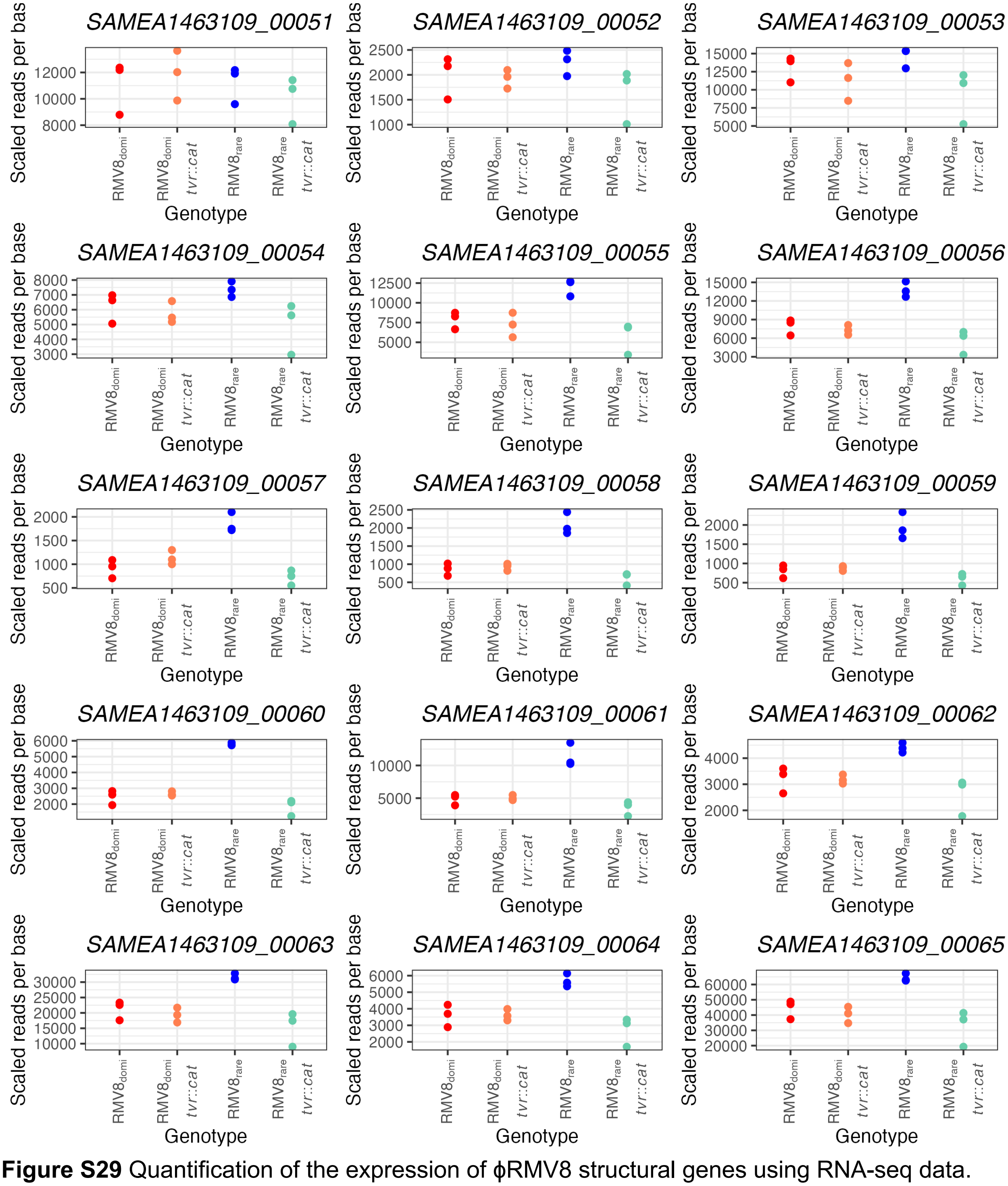


**Figure S29** Quantification of the expression of ϕRMV8 structural genes using RNA-seq data. Points are coloured using the scheme described in Fig. 4. Each plot shows the transcription of a different gene, in scaled reads per base, across the four genotypes. These genes tend to be most highly expressed in $RMV8_{rare}$.

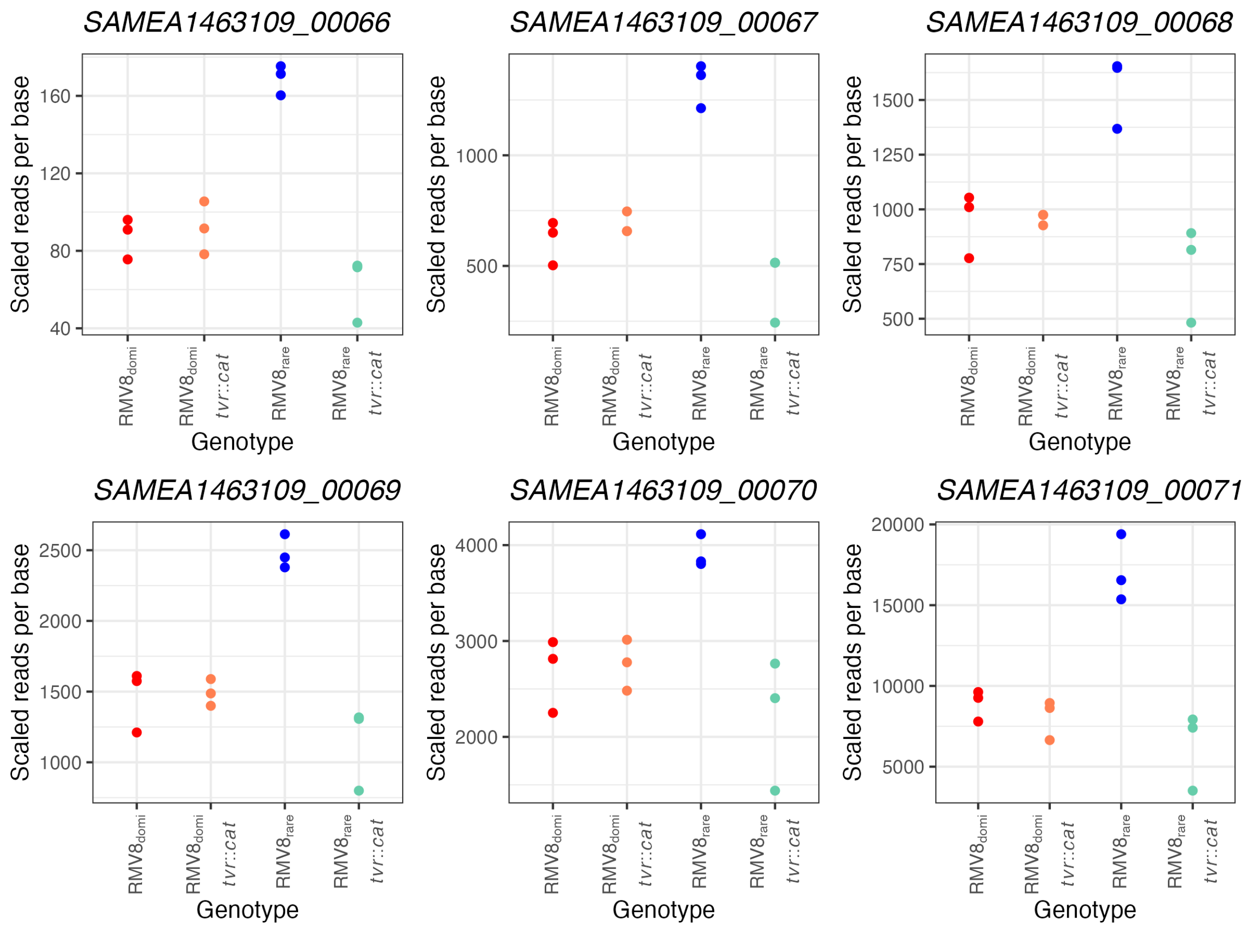


**Figure S30** Quantification of the expression of ϕRMV8 lysis genes using RNA-seq data. Points are coloured using the scheme described in Fig. 4. Each plot shows the transcription of a different gene, in scaled reads per base, across the four genotypes. These genes tend to be most highly expressed in $RMV8_{rare}$.

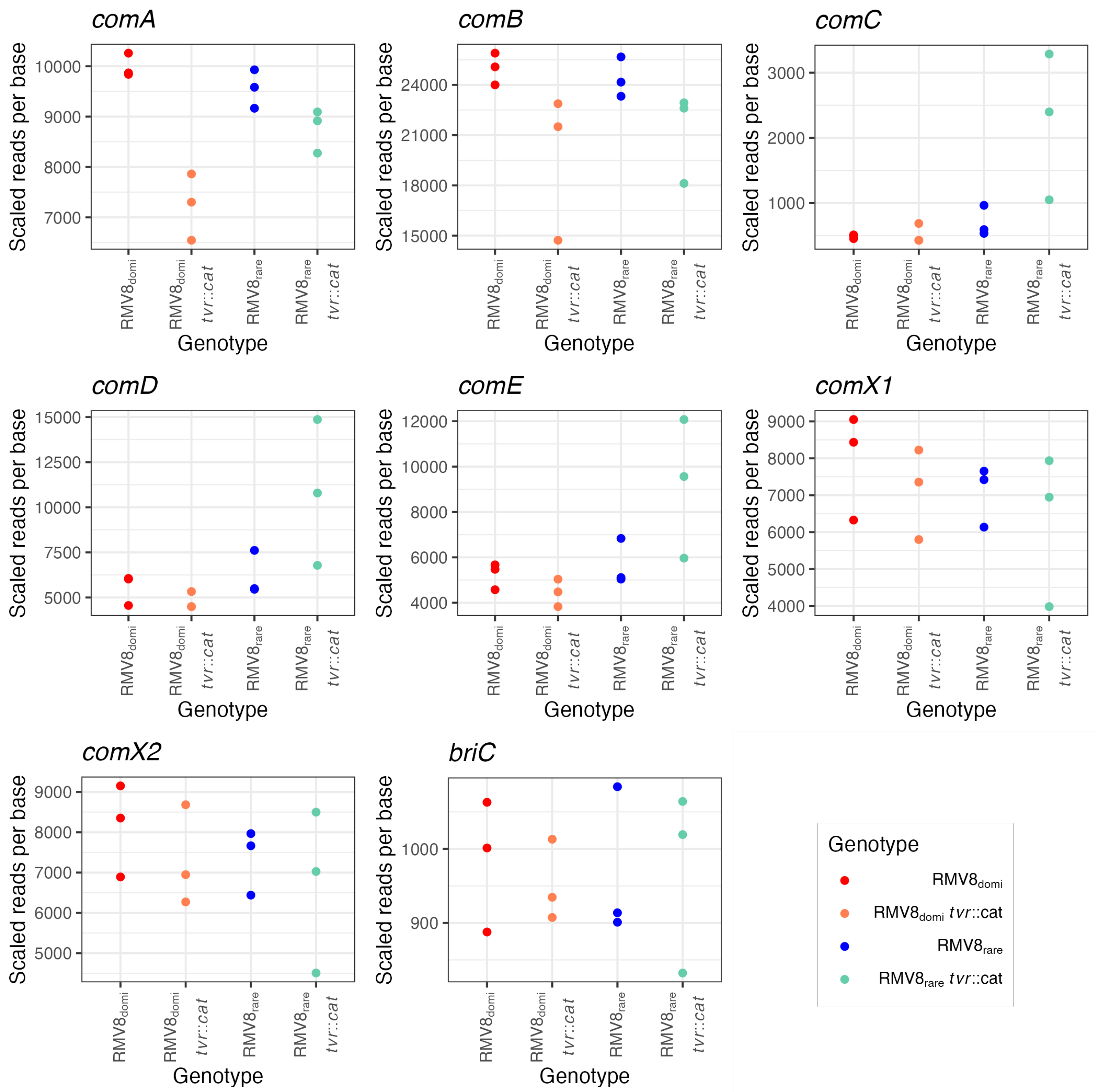


**Figure S31** Quantification of the expression of early competence genes using RNA-seq data. Points are coloured using the scheme described in Fig. 4, as shown in the key. Each plot shows the transcription of a different gene, in scaled reads per base, across the four genotypes. The *comCDE* operon was more highly transcribed in $RMV8_{rare}$ *tvr::cat* than the other genotypes, although this difference was only significant for *comC*.

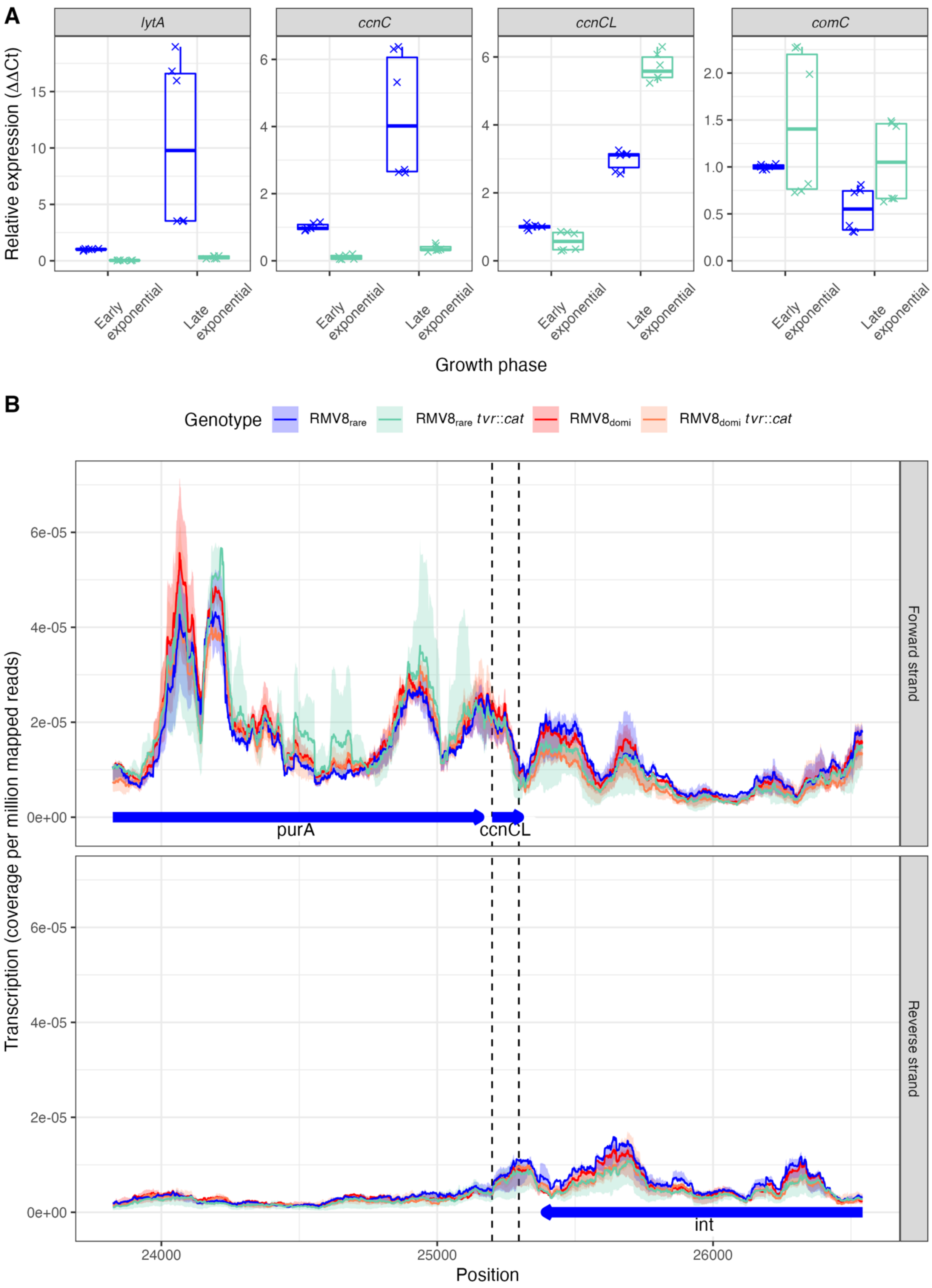
A
lytA
ccnC
ccnCL
comC
Relative expression (ΔΔCt)
Early exponential
Late exponential
Growth phase
B
Genotype
RMV8rare
RMV8rare tvr::cat
RMV8domi
RMV8domi tvr::cat
Forward strand
Reverse strand
purA
ccnCL
int
Transcription (coverage per million mapped reads)
Position

**Figure S32** Effects of phage activity on the expression of *ccnC*, *ccnCL* and *comC*. (A) Comparison of gene expression in $RMV8_{rare}$ and $RMV8_{rare}$ *tvr*::*cat* during the early ($OD_{600}$ = 0.2) and late ($OD_{600}$ = 0.5) exponential phase using qRT-PCR. Levels of transcription were quantified relative using the ΔΔCt approach (see Methods). Each point represents to one of six measurements per gene, corresponding to three technical replicate measurements of each of two biological replicates. The interquartile range and median are summarised by the boxplots. (B) RNA-seq data at the ϕRMV8 *attL* site. The line graphs show the RNA-seq coverage, in reads per million reads mapped, of each base in the displayed *attL* region of the *S. pneumoniae* $RMV8_{rare}$ genome. The lines show the median of three replicates, and are coloured according to the genotype from which they arose. The full range of the replicates across all bases is shown by the shaded ribbon. The top panel shows transcription of the forward strand, with the blue arrows indicating the corresponding genes transcribed in this direction. The bottom panel shows the equivalent data for the reverse strand of the genome. The vertical dashed lines show the boundaries of the *ccnCL* gene. Quantification of the sense transcription of this locus on the forward strand by qRT-PCR is likely to be distorted by antisense transcription driven by the prophage's lysogeny genes on the reverse strand.

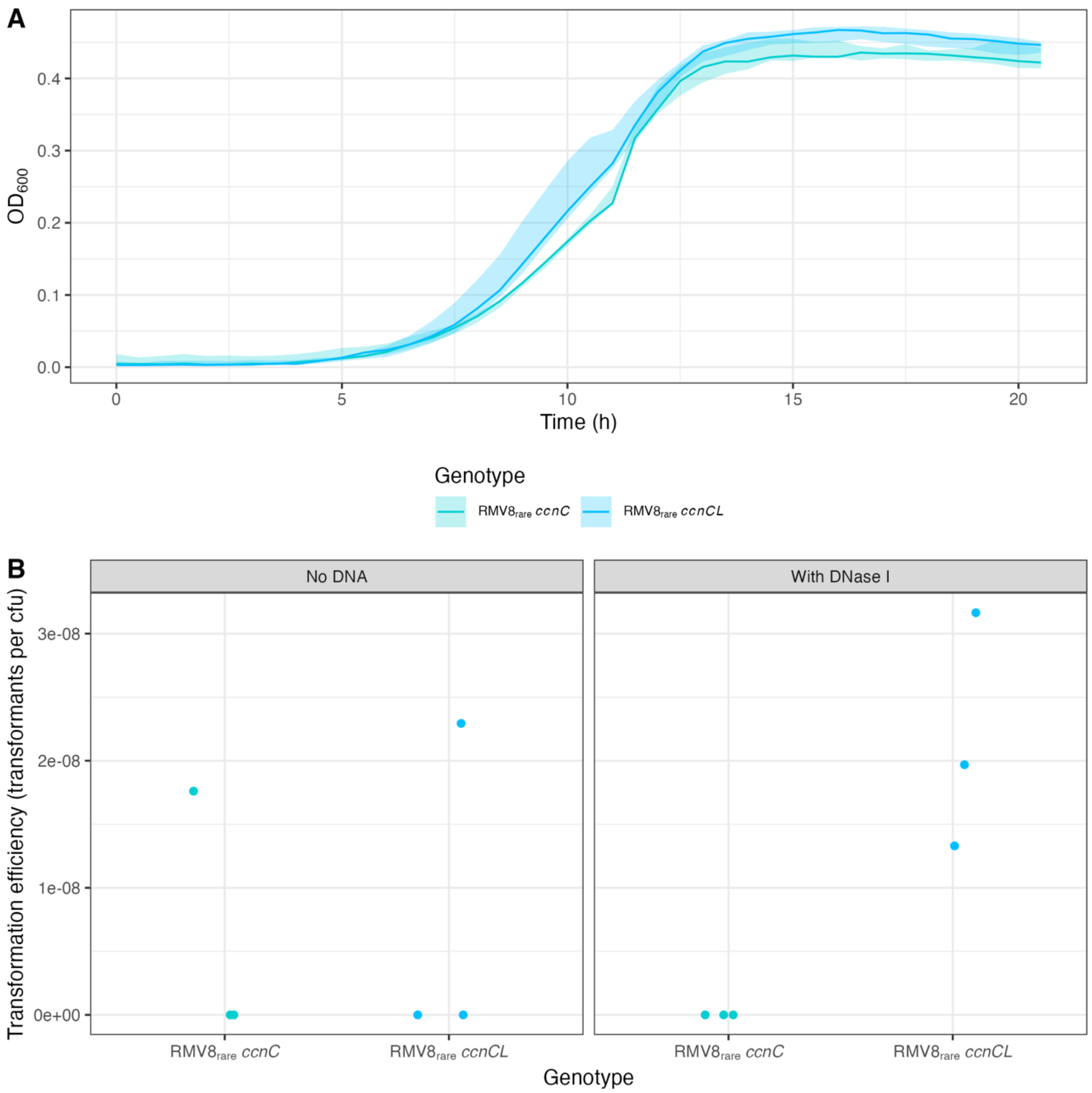


**Figure S33** Characterisation of RMV8$_{rare}$ *ccnC* and *ccnCL* mutants. (A) Growth curves comparing RMV8$_{rare}$ *ccnC* and *ccnCL* mutants using three replicates. The solid line represents the median, and the ribbon shows the range between the minimum and maximum. (B) Negative controls for transformation experiments with RMV8$_{rare}$ mutants. These experiments were designed to test whether the rifampicin-resistant colonies isolated in the absence of added CSP were arising through transformation. In one set of experiments, the protocol was undertaken with no exogenous DNA. A maximum of one rifampicin-resistant colony was isolated in each experiment, likely representing the infrequent emergence of this resistance phenotype through spontaneous mutation. Similarly, the protocol was undertaken with exogenous DNA and DNase I. This again yielded few rifampicin-resistant colonies, demonstrating that the rifampicin-resistant colonies observed in the transformation experiments were primarily generated through uptake of exogenous naked DNA.

### Supplementary Tables

All tabulated data are available in the associated spreadsheet.

**Table S1** Accession codes and epidemiological data for the 891 GPSC12 isolates analysed in Fig. 1.

**Table S2** Statistics on the diversification of Clades I-VI within GPSC12 calculated from the evolutionary reconstruction by Gubbins.

**Table S3** Distribution and sequences of the types of csRNAs identified in this work.

**Table S4** Accession codes of the RNA-seq datasets.

**Table S5** Statistical analysis of gene expression from RNA-seq data. All genetic features are labelled with their locus tags in the annotation of $RMV8_{rare}$ (accession code OX244288). Each row describes the output for an individual gene in a single RNA-seq dataset.

**Table S6** Significant changes in expression identified by RNA-seq analyses. All the listed genes differed in expression between genotypes with a *q* value exceeding the threshold value of $10^{-3}$.

**Table S7** Genotypes used in experiments described in this study.

**Table S8** Oligonucleotides used in experiments described in this study.
